# Microsecond dynamics control the HIV-1 envelope conformation

**DOI:** 10.1101/2023.05.17.541130

**Authors:** Ashley L. Bennett, R.J. Edwards, Irina Kosheleva, Carrie Saunders, Yishak Bililign, Ashliegh Williams, Katayoun Manosouri, Kevin O. Saunders, Barton F. Haynes, Priyamvada Acharya, Rory Henderson

## Abstract

The HIV-1 Envelope (Env) glycoprotein facilitates host cell fusion through a complex series of receptor-induced structural changes. Although significant progress has been made in understanding the structures of various Env conformations and transition intermediates that occur within the millisecond timescale, faster transitions in the microsecond timescale have not yet been observed. In this study, we employed time-resolved, temperature-jump small angle X-ray scattering to monitor structural rearrangements in an HIV-1 Env ectodomain construct with microsecond precision. We detected a transition correlated with Env opening that occurs in the hundreds of microseconds range and another more rapid transition that preceded this opening. Model fitting indicated that the early rapid transition involved an order-to-disorder transition in the trimer apex loop contacts, suggesting that conventional conformation-locking design strategies that target the allosteric machinery may be ineffective in preventing this movement. Utilizing this information, we engineered an envelope that locks the apex loop contacts to the adjacent protomer. This modification resulted in significant angle-of-approach shifts in the interaction of a neutralizing antibody. Our findings imply that blocking the intermediate state could be crucial for inducing antibodies with the appropriate bound state orientation through vaccination.

## Introduction

The HIV-1 envelope glycoprotein (Env) mediates viral attachment and entry into host cells (*1, 2*). This is accomplished through a complex series of receptor CD4 and co-receptor CCR5/CXCR4 induced Env structural transitions that ultimately eventuate in virion-host cell membrane fusion and host cell infection (*3-5*). The Env ectodomain is composed of three hetero-protomers, each consisting of a receptor-binding gp120 domain and fusion machinery containing the gp41 domain (Fig. 1A) (*1-4, 6*). In the prefusion closed state, the gp120 domains surround the gp41 domains, thus sequestering the fusion machinery prior to CD4 receptor and CCR5/CXCR4 co-receptor binding (*4*). Receptor CD4 engagement induces internal rearrangements and rotation of the gp120 domains to expose the gp41 fusion elements (*3, 4*). Env ectodomain structures determined using X-ray crystallography and cryo-electron microscopy (cryo-EM) have identified key structural elements within these domains that control these movements (*2, 4, 7-66*). Inter-protomer contacts between gp120 subunits at the membrane distant trimer apex and contact between each gp120 with a trimeric gp41 three-helix bundle characterize the closed state (Fig. 1B, left) (*67*). These stabilizing inter-and intra-protomer contacts are disrupted in the CD4-induced, open Env structure, resulting in an open state configuration (Fig. 1B, right) (*67*). Although these studies have provided detailed end-state configurations for this important transition, less is known about the structural intermediates the Env visits along this opening path.

**Figure 1:**
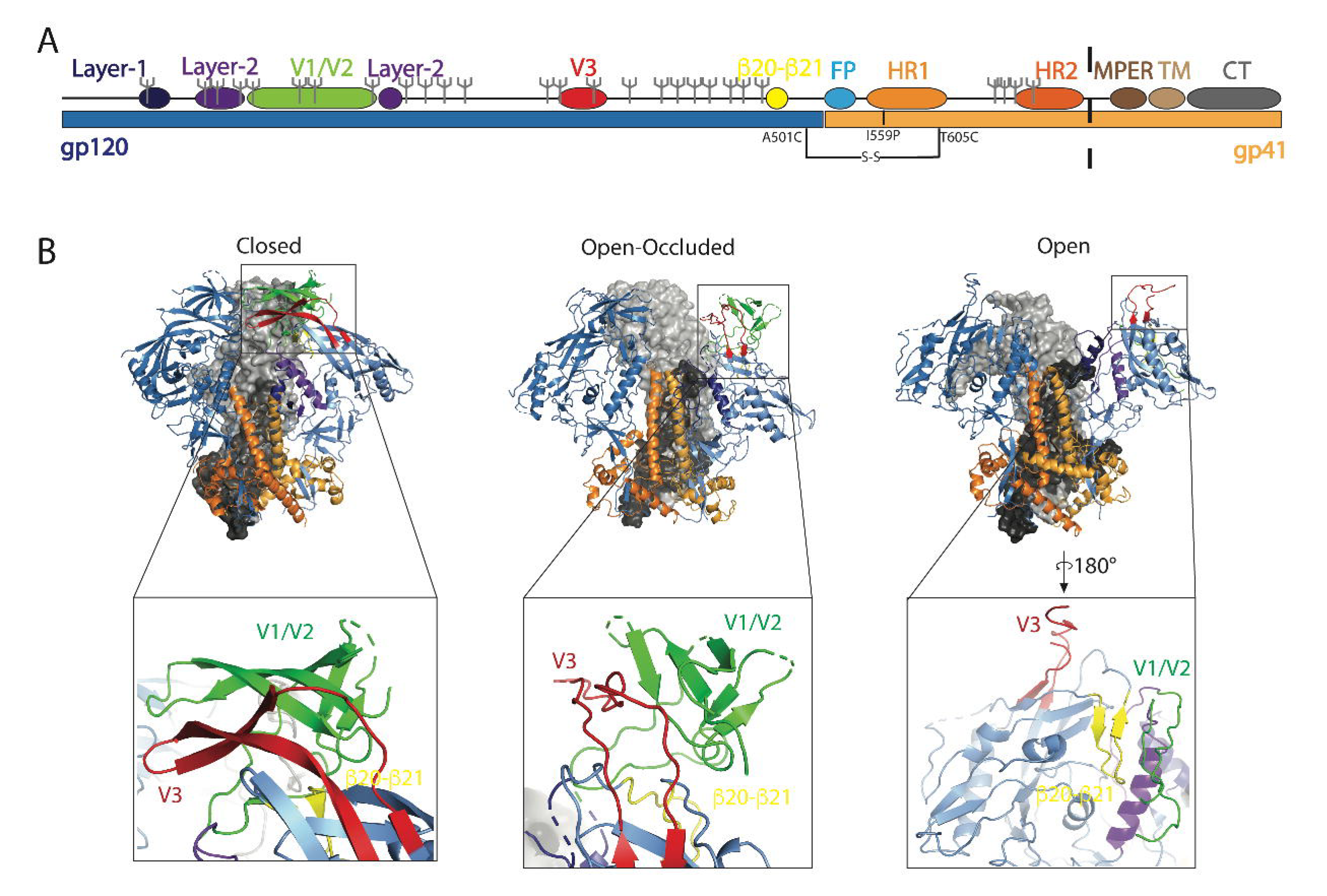
The HIV-1 Envelope Glycoprotein is Structurally Dynamic. **(A)** Linear sequence of the HIV-1 Env with gp120 in blue and gp41 in light orange and the layer-1, layer-2, variable domains 1 and 2 (V1/V2), variable domain 3 (V3), b20-b21, fusion peptide (FP), heptad repeat 1 (HR1), heptad repeat 2 (HR2), membrane proximal external region (MPER), transmembrane domain (TM) and the cytoplasmic tail (CT) allosteric elements are colored in dark blue, purple, green, red, yellow, sky blue, orange, dark orange, brown, tan, and grey, respectively. Glycosylation sites as predicted by Glycosite for CH505 models are denoted by grey forks. The black dashed line represents the location of truncation in SOSIP constructs. **(B)** Cryo-EM structures of a closed Env trimer (left) and an occluded Env trimer (middle) and an open Env trimer (right) from the side viewpoint. Allosteric elements of the gp120 and gp41 domains are colored identically to panel A.

Receptor-induced movement from the prefusion, closed state results in the elimination of apex inter-protomer contacts between two sequence variable loop containing segments, V1/V2 and V3 (*4, 34, 57, 58*). The final state for an Env ectodomain bound to CD4 and a co-receptor mimicking antibody consisted of rotated gp120s, an exposed, flexible V3 loop, and a dynamic, conformationally displaced V1/V2 (*30, 34*). Another structure determined with an Env ectodomain interacting with a broadly neutralizing antibody, b12, showed a similar rotation of gp120 but no major rearrangements in V1/V2 or V3 (*10*). Several additional antibody-bound structures with these gp120 rotated, V1/V2 occluded (open-occluded) configurations (Fig. 1B, middle) have been identified suggesting this state may be an intermediate in the closed to open transition (*10, 43, 44*). Single-molecule Förster resonance energy transfer (smFRET) studies and double electron-electron resonance (DEER) experiments (*68*) indicate that Env ectodomain is structurally dynamic in the absence of receptor (*69-72*). Correlation between these observations and known structures remains challenging and observable timescales of these measurements limit interrogation of rapid structural conversions that may control these movements.

Knowledge of Env structure and dynamics has nevertheless played an important role in HIV-1 vaccine immunogen design. Preventing transitions from the prefusion, closed configuration is thought to be essential for selecting functional improbable broadly neutralizing antibody (bnAb) mutations required for neutralization induction by vaccination (*2, 53, 54*). The Env glycoprotein is the primary target for these design efforts involving both full-length membrane-associated gp160 and truncated soluble gp140 constructs. The soluble ectodomains are often stabilized using an intra-protomer gp120 to gp41 linking disulfide (SOS) mutation and Ile 559→Pro (IP) mutation to prevent gp120 shedding and movements from the prefusion state, respectively (*2, 54*). Additional stabilization for these SOSIP immunogens has proven essential to ensure gp41 fusion elements are not exposed (*7, 18, 20, 25, 49, 53, 54, 63, 73-79*). These include structure-based design modifications within the allosteric components of gp120 to gp41 intra-and inter-protomer contacts toward the trimer base. Despite this additional stabilization, vaccination can still induce fully open state or open-occluded state targeting antibodies (*37, 43*). While smFRET experiments have identified intermediates in the Env opening pathway that occur in sub-millisecond timescales (*69-72*), these observations leave open the possibility that microsecond timescale structural movements intermediate to the already observed states have not been identified. An improved understanding of the Env opening pathway will provide useful information to guide the design of greater closed-state Env trimer stability.

Here, to probe Env opening on microsecond timescales, we performed time-resolved, temperature-jump (TR, T-Jump) small angle X-ray scattering (SAXS) experiments using a CH505 isolate (*80*) trimeric Env ectodomain SOSIP. These data revealed the rapid formation of a structural intermediate after ∼13 µs that moved to a distinct structural state after ∼700 µs. Structure-based modeling of these transitions suggested that the structural states correspond to the breaking of inter-protomer gp120 contacts at the trimer apex followed by gp120 domain rotation outward, away from the central axis. This led us to design an inter-protomer Env apex stapled construct that was designed to eliminate access to the first, rapidly forming intermediate. Structures of this stabilized construct complexed with receptor binding site-targeting antibodies revealed a closed Env state with one antibody (b12) showing marked shifts in its angles of approach compared to antibody binding to unstabilized Env. These results show that knowledge of transient intermediate structural states can guide vaccine immunogen design.

## Methods

### Recombinant HIV-1 envelope SOSIP production

A chimeric CH505 transmitted founder (CH505) envelope SOSIP, containing the CH505 gp120 and truncated BG505 gp41 domains, and a BG505 envelope SOSIP were expressed in Freestyle^TM^ 293-F cells (ThermoFisher Cat No. R79007). Before transfection, cells were diluted in Freestyle^TM^ 293 Expression Medium (Cat No. 12338018) to 1.25×10^6^ cells/mL at a volume of 950 mL. Plasmid DNA expressing the envelope SOSIP and furin were co-transfected at a 4:1 ratio (650 μg and 150 μg per transfection liter, respectively) and incubated with 293fectin^TM^ transfection reagent (ThermoFisher Cat No. 12347019) in Opti-MEM I Reduced Serum Medium (ThermoFisher Cat No. 31985062) to allow for complex formation. The diluted mixture was added to the cell culture which was incubated at 37°C, 9% CO_2_ on a shaker at 120 rpm for 6 days. On day 6 the cell supernatant was harvested by centrifuging the cell culture at 4000 xg for 45 minutes. The supernatant was filtered with a 0.45 μm PES filter and concentrated to approximately 100 mL using a Vivaflow^®^ 200 cross-flow cassette (Sartorius Cat No. VF20P2).

Envelope SOSIPs were purified using a PGT145 affinity chromatography column equilibrated in 15 mM HEPES, and 150 mM NaCl (pH = 7.1). A PGT145 Ab affinity column was made by coupling PGT145 mAbs to CNBr-activated Sepharose 4B (Cat No. 170430-01, GE Bio-Sciences) and packed into a Tricorn column (GE Healthcare). The supernatant was applied over the column at 2 mL/min using an AKTA go chromatography system (Cytiva) followed by three column volume wash steps with 15 mM HEPES, and 150 mM NaCl. Protein was eluted off the column using 3 M MgCl_2_ and diluted in 15 mM HEPES, and 150 mM NaCl buffer. The protein sample was buffer exchanged into 15 mM HEPES, and 150 mM NaCl by ultrafiltration using a 100 kDa MWCO Amicon^®^ Ultra-15 Centrifugal Filter Unit (Millipore Aldrich Cat No. UFC9010) and concentrated to <0.5 mL for size exclusion chromatography. Size exclusion chromatography was performed using a Superose 6 10/300GL Column (Cytiva) on an AKTA go system in 15 mM HEPES, and 150 mM NaCl. Fractions containing trimeric SOSIP were collected.

All lots produced were subjected to quality control including analytical SEC, SDS-PAGE, thermal shift analysis, biolayer interferometry (BLI), and negative stain electron microscopy (NSEM) to assure the presence of well-folded Env trimers. Production lots deviating from mean observed values in each experiment greater than one standard deviation were not used in the experiments. For the TR, T-jump SAXS experiments, all CH505 SOSIP lots were combined. The combined sample was subjected to the same quality control after completion of the TR, T-jump SAXS experiments to verify sample integrity.

### Thermal shift assay

Thermal shift assay was performed using Tycho NT.6 (NanoTemper Technologies). Envelope ectodomain SOSIPs were diluted (0.15 mg ml^−1^) in 15 mM HEPES buffer with 150 mM NaCl at pH 7.1. Intrinsic fluorescence was recorded at 330 nm and 350 nm while heating the sample from 35 to 95°C at a rate of 3°C min^−1^. The ratio of fluorescence (350/330 nm) and the inflection temperatures (Ti) were calculated by Tycho NT.6.

### Biolayer Interferometry

Sample mAb and Fab binding were obtained using biolayer interferometry (BLI; OctetR4, FortéBio). Antibodies were immobilized on anti-Human IgG Fc capture (FortéBio) sensor tips while Fabs were immobilized on Protein A sensor tips, each via immersion in 4.5 μg/ml mAb in 15mM HEPES buffer with 150 mM NaCl at pH 7.1 for 300 s followed by washing in 15 mM HEPES buffer with 150 mM NaCl at pH 7.1 for 60 s at 1000 rpm. The sensor tips were then immersed in the 45 μg/ml SOSIP-containing wells for 180 s. Reported binding corresponds to values and the end of the association phase. Data were evaluated using the Octet Data Analysis software (FortéBio).

### Negative-stain electron microscopy

An aliquot from RT was diluted to 20 µg/ml with 0.1% w/v n-Dodecyl β-D-maltoside (DDM) (50 mM Tris pH 7.4, 150 mM NaCl, 0.03 mg/mL sodium deoxycholate). The sample was immediately applied to a glow-discharged carbon-coated EM grid for 8-10 seconds, then blotted, and stained with 2 g/dL uranyl formate for 1 min, blotted, and air-dried. Grids were examined on a Philips EM420 electron microscope operating at 120 kV and nominal magnification of 49,000x, and 114 images were collected on a 76 Mpix CCD camera at 2.4 Å/pixel. Images were analyzed by 2D class averages using standard protocols with Relion 3.0 (*81*).

### Time-resolved, temperature-jump small angle X-ray scattering beamline setup

The BioCARS 14-ID-B beamline at the Advanced Photon Source at Argonne National Laboratory was used to conduct TR, T-Jump SAXS, and static temperature series SAXS experiments. CH505 protein was concentrated to a final concentration of 5.125 mg/mL in 15 mM HEPES buffer with 150 mM NaCl at pH 7.1, for a total volume of ∼4 mL. The protein sample was syringe filtered with a 0.22-micron filter and degassed before use. A peristaltic pump injected protein/buffer solution through a 700-micron quartz capillary mounted on a custom, temperature-controlled aluminum nitride (AlN) holder. Temperature jumps of the sample were initiated using a 7 ns infrared (IR) laser pulse at 1.443 micrometers wavelength and 1.1 mJ/pulse to excite the O-H stretch of water. The sample was probed with a 1.6 µs pulse of pink X-ray beam with photon energy 12 keV and bandpass of 300 eV after a variable time-delay (Supp. Fig. 3A). Images were collected using a Rayonix MX340-HS X-ray detector in scattering vector (q) range 0-2.5 Å^-1^.

### Temperature jump calibration

We measured the static SAXS signal of water at 3°C intervals from 25°C to 51°C. We then calculated static SAXS difference curves for Δ5°C, Δ8°C, and Δ11°C by subtracting the SAXS signals at 45°C and 40°C, 48°C and 40°C, and 51°C and 40°C, respectively (Supp. Fig. 4A). We then measured the TR, T-Jump SAXS signal of water at 10 µs post heating, starting at 42°C. The temperature jump was determined to be Δ6°C by linear regression analysis of the water TR, T-Jump and static SAXS difference signal intensity at q=2.3 Å^-1^ (Supp. Fig. 4B).

### Static SAXS

Static SAXS scattering profiles of BG505 (*82*) and the chimeric CH505 (*80*) Env SOSIPs were collected at the Lawrence Berkely National Laboratory Advanced Light Source SIBYLS 12.3.1 beamline (*83-86*). Samples were shipped overnight at −79°C and stored at −80°C before data collection. A series of concentrations between 0.5 mg/ml and 5 mg/ml were collected with a total exposure time of 10 s. Data were reduced and analyzed using the ATSAS program suite.

### Static SAXS temperature series

Static SAXS temperature series data were measured at the Advanced Photon Source BioCARS 14-ID-B beamline used for TR, T-Jump SAXS, described above. both 15 mM HEPES buffer and 5.125 mg/mL CH505 Env sample at 25°C, 35°C, 44°C, and 50°C (Supp. Fig. 2A). Static SAXS signals were collected using 24 bunches in continuous translation mode at 20Hz on the BioCARS 14-ID-B beamline. A total of 50 images were collected for each set for scattering vector range q=0-2.5 Å^-1^.

### CH505 Time Delay Series

Time-resolved, temperature-Jump SAXS data were collected at the Advanced Photon Source BioCARS 14-ID-B beamline. Data were collected for a 5.125 mg/mL CH505 Env SOSIP sample at several different time delays after sample heating from 44°C to ∼50°C by IR pulse temperature-jump: 500 ns, 1.5 µs, 3.0 µs, 5.0 µs, 10 µs, 50 µs, 100 µs, 500 µs, 1 ms, 10 ms, and 100 ms. For each time delay measured, TR, T-Jump SAXS was also measured at −10 µs and −5 µs prior to heating with IR laser (‘laser off’; Supp. Fig. 3A). To accommodate the large change in timescales and minimize systematic errors due to experimental drift, we measured TR, T-Jump SAXS for buffer and Env in multiple sets, with overlapping time delays between each set. The CH505 Env SOSIP TR, T-Jump SAXS was measured in three sets. Env set 1 includes −10 µs, −5 µs, 10 µs, 50 µs, 100 µs, 500 µs, and 1 ms measured with 24 bunches in continuous translation mode at 20 Hz (*87*), collecting 250 images for each time delay. Set 2 for CH505 Env SOSIP was measured with 11 bunches in 20 Hz continuous translation mode (*87*) and includes −10 µs, −5 µs, 1.5 µs, 3 µs, and 5 µs, with 362 images for each time delay. 200 images were collected for Env set 3, which was measured in 5 Hz step translation mode with 24 bunches and included time delays −10 µs, −5 µs, 1 ms, 10 ms, and 100 ms. 25 images for the 500 ns time-delay were collected separately using a single bunch at 20 Hz continuous translation mode (*87*) with the T-Jump starting from 42°C.

TR, T-Jump SAXS profiles of buffer were collected using the same protocol. For buffer set 1, 100 images each at −10 µs, −5 µs, 1.5 µs, 3.0 µs, and 5.0 µs time delays were collected with 11 bunches in 20 Hz continuous translation mode (*87*). Buffer set 2 was collected with 24 bunches in 20 Hz continuous translation mode (*87*) and consists of 50 images each at time delays −10 µs, −5 µs, 5 µs, 10 µs, 25 µs, 50 µs, 100 µs, 250 µs, 500 µs, and 1 ms. HEPES set 3 was measured in 25 bunches in 5 Hz step mode (*87*), collecting 50 images each at −10 µs, −5 µs, 1 ms, 10 ms, and 100 ms time delays.

### Data Reduction

Scattering intensity (I) was binned as a function of the scattering vector (q, Å^-1^) and radially averaged to produce isotropic scattering curves of the scattering vector, q, and intensity at q, I(q), with q calculated according to Eq. 1:

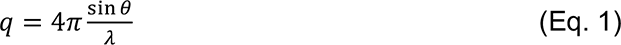

where ***θ*** is one half the scattering angle and λ is the X-ray wavelength. A mask was applied below 0.02 Å^-1^ q to eliminate the scattering signal near the beam stop for all collected images for static and TR, T-Jump SAXS data sets collected on the BioCARS 14-ID-B beamline. The scattering curves were normalized to the isosbestic point for water, 1.4-1.6Å^-1^. This data reduction was performed with custom software at the BioCARS beamline.

### Data Processing

For static SAXS scattering curves, outliers were detected using two iterations of singular value decomposition (SVD) to remove curves with first right vector values exceeding 2.5 standard deviations from the mean. Once outliers were removed, average curves for each temperature were calculated for both buffer and CH505 Env SOSIP samples. The procedure was identical for both buffer and CH505 Env SOSIP. The remaining curves were used to determine the average static curves for buffer and CH505 Env SOSIP. The average buffer curves were scaled to the average CH505 curves in the scattering vector range q=1.5-2.5 Å^-1^, with the scaling factors (s) calculated according to Eq. 2:

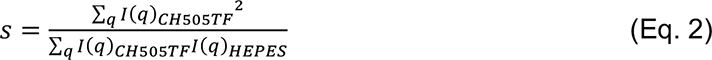

The scaled average buffer scattering curves were subtracted from the average CH505 static scattering curves for each temperature to produce buffer-corrected average static SAXS scattering curves.

To identify outliers for TR, T-Jump SAXS curves, two iterations of SVD for both laser on and laser off scattering curves were used to detect scattering curves with first right vector values exceeding 2.5 standard deviations from the mean. A second iteration of SVD analysis was performed to ensure that any outliers detected in the first SVD iteration did not skew the calculation of the mean. This outlier analysis was carried out for −10 µs and −5 µs laser off scattering profiles as well as the scattering profiles for each time delay after heating with the IR laser pulse. Difference curves were then calculated for the remaining scattering curves by subtracting the −10 µs laser-off curve from each time delay. An iterative *χ*^2^ analysis was used to detect difference curves with a *χ*^2^ > 1.5 for TR, T-Jump SAXS at each time delay. The same procedure was performed on both buffer and CH505 Env SOSIP. Average difference curves for each time delay were calculated from the remaining difference curves. For each time delay difference profile, the corresponding average buffer difference curves were scaled using Eq. 2 to the average CH505 Env difference curves in the buffer scattering region (q=1.5-2.5 Å^-1^) and subtracted from the respective average CH505 difference profiles to yield average, buffer-corrected TR, T-Jump SAXS difference curves for CH505 Env SOSIP. These buffer-subtracted curves were used for the remaining TR, T-Jump and static SAXS data analysis.

Average buffer-subtracted CH505 scattering profiles were determined using the laser-on scattering curves for CH505 Env SOSIP and buffer T-Jump scattering curves. The same procedure was followed for determining these buffer-subtracted T-Jump scattering curves as for the static SAXS curve processing described above. Guinier analysis was performed in ATSAS primus software (*88*) to determine the particle radius of gyration (R_g_) and to inspect for radiation damage and aggregation for both static (Supp. Fig. 1C, D; Supp. Fig. 2C-F) and TR, T-Jump SAXS (Table 1) scattering data. Scattering profile Kratky plots and pair-distance distributions (P(r)) were also assessed for each static (Supp. Fig. 1B, E; Supp. Fig. 2B, G) and TR, T-Jump (Supp. Fig. 3C, D) scattering profiles in ATSAS.

**Table 1:** Radius of gyration (R_g_) values determined for experimental TR, T-Jump SAXS scattering profiles.

| Table 1: Experimental $R_g$ | | | |
| --- | --- | --- | --- |
| Data Set | Guinier $R_g$ (Å) | P(r) $R_g$ (Å) | dmax (Å) |
| 500ns | 42.98 | 50.22 | 145.66 |
| 1.5us | 46.14 | 51.19 | 154.63 |
| 3us | 46.1 | 51.19 | 154.5 |
| 5us | 46.12 | 51.2 | 154.53 |
| 10us | 42.76 | 51.07 | 152.59 |
| 50us | 42.71 | 51.05 | 153.46 |
| 100us | 42.74 | 51.09 | 152.49 |
| 500us | 42.79 | 51.06 | 153.24 |
| 1ms | 42.79 | 51.13 | 153.23 |
| 10ms | 40.79 | 51.25 | 153.85 |
| 100ms | 40.77 | 51.22 | 154.5 |

### TR, T-Jump Component Analysis

A singular value decomposition (SVD) based analysis of the averaged, buffer-subtracted CH505 TR, T-Jump difference curves, and CH505 static temperature series scattering curves was conducted to decompose the SAXS signals. The singular values were used to determine that there are two component signals (Supp. Fig. 5A). The SVD left vectors can reveal individual feature curves contributing to the SAXS profiles while the right vectors show the relative contributions of the corresponding left vectors at each time delay or temperature. Individual components from the buffer subtracted CH505 Env SOSIP temperature series static SAXS curves were also identically extracted by SVD (supp. Fig. 5E, F).

### TR, T-Jump Kinetic Analysis

Kinetic analysis was carried out using a bootstrapping method using 100 resamples from the CH505 TR, T-Jump difference curves. For each time delay in the CH505 TR, T-Jump data set, 80% of the difference curves remaining after outlier subtraction were selected and averaged. The previously determined average TR, T-Jump buffer curves were subtracted from the respective bootstrapped TR, T-Jump CH505 curve to yield to the buffer-corrected, bootstrapped average curves for each time delay. The area under the bootstrapped difference curve (AUC) for each CH505 Env time delay was determined according to Simpson’s rule in the q range 0.02-0.1Å^-1^. The AUC vs. time delay plots were fit to a double exponential with the form shown in Eq. 3, as indicated by the SVD singular values.

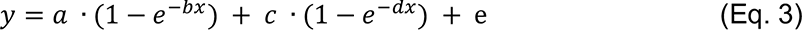

where *b*=1/t_fast_ and d=1/t_slow_.

After bootstrapping, the first two right vectors and the AUC were fit to the double exponential decay function in Eq. 3 to extract kinetic parameters for the individual CH505 TR, T-Jump scattering components.

### Modeling

Atomic models of CH505 Env in closed (Closed), open-occluded (OpenOccluded), and open (Open) conformations were built using Modeller10.1 (*89*). The templates used for the models included PDB ID 6UM6 (*90*) for closed protomers, PDB ID 6CM3 (*30*) for the open protomers, and PDB ID 7TFN (*43*) for the open-occluded protomers. Sequence alignments between CH505 Env SOSIP and templates were conducted using Clustal Omega in the DNASTAR MegAlign software. Disulfide patches were added to Modeller scripts to ensure the formation of disulfide bonds.

Glycosite (*91*) was used to predict the possible N-linked glycosylation sites in the CH505 Env and Glycosylator (*92*) was used to glycosylate each model with mannose-9 (Man9) glycans. Three rounds of refinement were performed to remove glycan clashes. All rounds of refinement used 10 iterations, a 0.01 mutation rate, and a population size of 30. The first two rounds of refinement used 10 generations, while the last refinement used 20 generations. The number of individues was decreased by two between each refinement.

The theoretical SAXS profiles for each glycosylated model were calculated using the FoXS server (*93*). P(r) distributions for these theoretical SAXS scattering profiles were determined in ATSAS primus software and ΔP(r) curves were calculated by subtracting the closed P(r) curve from the Open or OpenOccluded P(r) curve. (*88*) and compared to the experimental and REGALS SAXS profiles and ΔP(r) curves. SVD analysis was performed on the theoretical scattering curves for the Closed, OpenOccluded, and Open models both with mannose-9 glycans and without glycosylation to extract component signals for transitions between these structures. Guiniers analyses (Table 3) were performed on each theoretical SAXS profile to determine the R_g_ (Table 3).

**Table 2:** Kinetic fits to TR, T-Jump SAXS data.

| Kinetic Fits |  |  |  |  |  |  |  |
| --- | --- | --- | --- | --- | --- | --- | --- |
| Data Set | a | b | c | d | e | 1/b | 1/e |
| RV1 | $-0.223 \pm 0.001$ | $0.00197 \pm 0.00002$ | $-0.1056 \pm 0.0009$ | $0.251 \pm 0.004$ | $0.536 \pm 0.001$ | $507 \pm 6$ | $3.98 \pm 0.07$ |
| RV2 | $0.86 \pm 0.09$ | $0.0016 \pm 0.0004$ | $0.22 \pm 0.04$ | $0.09 \pm 0.04$ | $-0.6 \pm 0.1$ | $606 \pm 147$ | $12 \pm 5$ |
| AUC | $0.00008936 \pm 0.00000009$ | $0.0746 \pm 0.0002$ | $0.0002966 \pm 0.0000003$ | $0.001412 \pm 0.000003$ | $-0.0005231 \pm 0.0000003$ | $13.34 \pm 0.04$ | $708 \pm 2$ |

**Table 3:** Radius of gyration (R_g_) values determined for theoretical SAXS scattering profiles.

| **Table 3: Theoretical $R_g$** |  |  |  |
| --- | --- | --- | --- |
| **Model** | **Guinier**<br>**$R_g$ (Å)** | **P(r) $R_g$**<br>**(Å)** | **dmax (Å)** |
| Closed + Man9 | 52.11 | 51.97 | 153.56 |
| OpenOccluded + Man9 | 53.36 | 53.32 | 161.36 |
| Open + Man9 | 50.85 | 50.9 | 156.2 |

### Molecular Dynamics

The CHARMM-GUI Glycan Modeler (*94-99*) was used to glycosylate Glycosite-predicted glycosylation sites with mannose-5 glycans and prepare gp120 systems for simulation. A total of 250, 5 µs trajectories were run for closed gp120 at 50°C. The Amber22 software package with pmemd CUDA implementation (*100-102*) in the CHARMM36 force field (*103-106*) and TIP3P water model (*107*) was used for simulations. A 150 Å octahedral box was used to allow for V1/V2 and V3 dislocation during simulation. Both systems were neutralized with 150 mM NaCl. 10,000 steps of energy minimization were performed with 1000 steps of steepest descent minimization with 500 kcal/mol•Å restraints on protein. A second, unrestrained energy minimization consisting of 10,000 steps and 1000 steepest-descent steps followed. Minimization was followed by 20 ps of NVT heating with 10 kcal/mol•Å restraints applied to protein atoms and a subsequent 5 ns NPT equilibration with no restraints, with a constant temperature of 323.15 K maintained using a Langevin thermostat (*108*), and pressure of 1 atm was maintained with isotropic position scaling. 250 ns of unrestrained simulations in the NVT ensemble were performed to equilibrate the system, followed by 5 µs of production simulation in the NVT ensemble. Electrostatic interactions were calculated with the Particle Mesh Ewald method (*109*) with a cutoff of 12Å and switching distance of 10 Å. The SHAKE algorithm (*110*) with hydrogen mass repartitioning (*111*) was used to constrain hydrogen atoms and allow for a 4 fs timestep. The cpptraj tool in AmberTools21 (*112*) was used to determine the RMSD time series of V1/V2 (residues 129-133, 135-150, 154-163, 168-177, 181-183, and 190-193 in HXB2 isolate numbering), V3 (residues 297-308 and 311-319 in HXB2 isolate numbering), and gp120 core (residues 288-291, 280-284, 355-358, 370-374, 377-382, 411-416, 439-452, and 462-466 in HXB2 isolate numbering) b-sheet a-carbon for each replicate of MD simulation. A custom Python script was used to determine the average RMSD and frequency distributions for all replicates aggregated together.

### Rational structure-based design

The CH235.UCA bound CH505.M5.G458Y SOSIP trimer structure (6UDA) was prepared in Maestro (*113*) using the protein preparation wizard (*114*) followed by in silico mutagenesis using Schrödinger’s cysteine mutation (*115*) and residue scanning (*116*) tools. Residue scanning was first performed for V1/V2 and V3 region inter-protomer contact residues. Scores and visual inspection were used in the selection of the prepared construct. The selected design was prepared in a previously reported CH505 isolate SOSIP backbone (42) with the introduction of an expression-enhancing 2P mutation (112), an allosteric machinery disabling set of mutations, F14 (49), and a CD4bs antibody binding enhancing N197D mutation (*117*).

### Cryo-electron microscopy sample preparation, data collection, and processing

Purified Envelope SOSIP ectodomain preparations were prepared at concentrations of 4-5 mg/ml in 15 mM HEPES buffer with 150 mM NaCl at pH 7.1 and mixed with Fab at a 1:5 molar ratio. A total of 2.5 µl of the complex was deposited on a CF-1.2/1.3 grid that had been glow-discharged for 15 s in a PELCO easiGlow glow discharge cleaning system. After 30-s incubation in >95% humidity, the excess protein was blotted away for 2.5 s before being plunge-frozen in liquid ethane using a Leica EM GP2 plunge freezer (Leica Microsystems). Frozen grids were imaged in a Talos Arctica (Thermo Fisher) equipped with a K3 detector (Gatan). Individual frames were aligned, dose-weighted, and CTF corrected followed by particle picking, 2D classification, ab initio model generation, heterogeneous refinements, homogeneous 3D refinements, and local resolution calculations in cryoSPARC (Supp. Table 1) (113).

### Cryo-electron microscopy structure fitting and analysis

The CH505.M5.G458Y SOSIP structure (6UDA) and either CH235.12 (5F96) or b12 (2NY7) structures were used to fit the cryo-EM maps in ChimeraX (114, 115). Mutations were made in PyMol (*118*). Coordinates were then fitted using Isolde (116) followed by iterative refinement using Phenix (*119*) real-space refinement. Structure and map analyses were performed using PyMol, Chimera (117), and ChimeraX (114, 115).

## Results

### Small angle X-ray scattering captures conformational transitions in HIV-1 Env glycoproteins

Time-resolved, temperature-jump SAXS experiments are referred to as pump-probe experiments. In the pumping stage, the system is perturbed, in this case by rapid heating of the water surrounding the sample by an IR laser. The probe stage occurs at a time delay relative to the pump stage and acts as a readout for the state of the system (Supp. Fig. 3A), in this case in the form of a SAXS profile that reports on the structural state of the protein. Successful application of this method, therefore, requires a construct that can readily undergo conformational transitions with the applied temperature jump and where the conformational change can be detected by SAXS measurement. We, therefore, sought first to identify an Env construct suitable for TR, T-Jump SAXS experiments. We used biolayer interferometry (BLI) to measure conformation-specific antibody binding to two SOSIP stabilized Env gp140 ectodomains from BG505 (*82*) and a CH505 transmitted founder virus derived Env chimera (CH505; CH505 derived gp120 and a BG505 derived gp41) (*80*). Both BG505 and CH505 were isolated from infected individuals (*80, 82*). Binding was tested to antibodies PGT145 (Env closed state-specific, trimer apex targeting) (*9*), 17b (Env open state-specific, coreceptor binding site targeting) (*120*), and the 19b (Env open state-specific, linear V3 tip epitope targeting) (*121*). While CD4-induced Envs in the open conformation bind 17b and 19b efficiently, Envs from certain HIV-1 isolates access these open conformations and bind 17b and 19b even in the absence of CD4 (*69-72*). PGT145 binds a quaternary epitope contributed by all three protomers and its binding confirms the presence of a closed trimer (*69, 71*). Both CH505 and BG505 SOSIP bind PGT145 confirming the presence of closed trimers in both Env preparations (Fig. 2A, right). CH505 SOSIP interacts with both 17b (Fig. 2A, left) and 19b (Fig. 2A, middle) while BG505 does not, indicating that CH505 SOSIP is more structurally labile and may, therefore, more readily respond to perturbation in the temperature-jump experiments.

**Figure 2:**
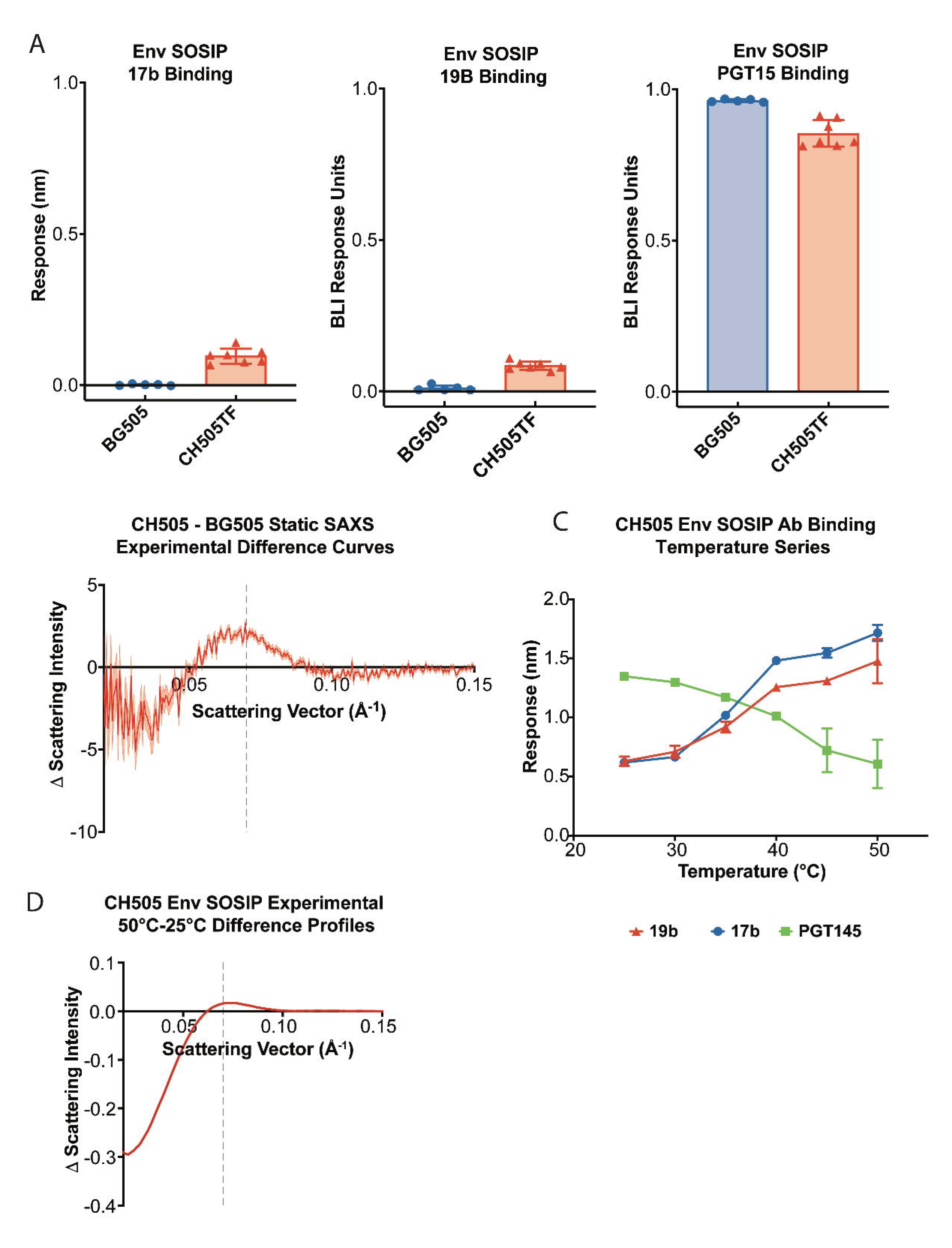
Static Small Angle X-Ray Scattering Profiles Capture HIV-1 Env Opening. **(A)** Biolayer interferometry (BLI) binding response for 17b (left), 19b (middle), and PGT145 (right) for BG505 SOSIP (blue circles) and CH505TF SOSIP (red triangles). The error bars indicate the standard deviation from the arithmetic mean aggregated for 2 experiments each with five replicates for BG505 and 7 replicates for CH505 Env SOSIPs. **(B)** Static SAXS scattering difference curve calculated by subtracting BG505 Env SOSIP SAXS scattering profile from the CH505 Env SOSIP with the propagated standard error (SE) shown as the shaded region. Scattering difference feature (q) peak at ∼0.07 Å^-1^ is indicated by the grey dashed line. **(C)** CH505 Env SOSIP temperature series binding response for interactions with 17b (blue circles), 19b (red triangles), and PGT145 (green squares). The error bars indicate the SD within the arithmetic mean for a total of three replicates for each measurement. **(D)** SAXS scattering difference curves calculated by subtracting CH505 Env SOSIP SAXS scattering profile at 25°C from the CH505 Env SOSIP scattering profile at 50°C. Scattering difference feature peak indicated at q = 0.07 Å^-1^ by the grey dashed line.

SAXS data for proteins collected in solution involve isotropic scattering due to molecular movement, as a result, the scattering curves report low-resolution features. Identifying specific curve features that correlate with differences in specific structural features of interest aid in assigning the observed transitions to structural changes. Owing to the stark differences exhibited by CH505 and BG505 SOSIPs in their binding to conformation-specific antibodies (Fig. 2A), we hypothesized that a clear differentiating feature would be found between their respective scattering profiles. We measured static SAXS profiles for CH505 and BG505 SOSIPs (Supp. Fig. 1A) and calculated difference profiles by subtracting the BG505 Env SOSIP scattering curve from the CH505 Env SOSIP scattering curve (Fig. 2B). The difference profiles show a noticeable feature with a peak at a scattering vector (q) value of 0.07 Å^-1^ with a downward trend between 0.01 and 0.05 Å^-1^ followed by a rise between 0.00 and 0.01 Å^-1^ (Fig. 2B). These results show that Env constructs that differ in their propensity to access open states exhibit differing SAXS profiles.

### HIV-1 Env glycoprotein conformational transitions can be induced by increasing temperature

The TR, T-jump SAXS experiments perturb the conformational equilibrium via a temperature jump initiated by a rapid IR laser pulse. It is therefore essential that the transition of interest occur within the temperature range sampled within the experiment. We selected CH505 SOSIP for these experiments due to its observed conformational lability and its ability to access the Env open state even in the absence of CD4 induction. To determine the effects of increasing temperature on the conformational dynamics of the CH505 SOSIP Env we measured 17b, 19b, and PGT145 binding at 25°C, 30°C, 35°C, 40°C, 45°C, and 50°C. As expected, elevated temperatures increased 17b and 19b binding while reducing PGT145 binding, demonstrating an increase in the population of open or open-like states at higher temperatures (Fig. 2C).

We next asked whether SAXS would capture structural changes in the CH505 SOSIP Env associated with increased temperature. We measured static SAXS scattering profiles for the CH505 SOSIP at 25°C, 35°C, 40°C, 44°C, and 50°C (Supp. Fig. 2A). The difference profiles display the same features as observed in the difference profiles for CH505 and BG505 SOSIPs between q=0.01-0.05 Å^-1^ and at 0.07 Å^-1^ (Fig. 2D), providing further evidence that the observed differences between BG505 and CH505 Envs occurred due to their differing propensity to access the open states. Unlike the static difference profile between CH505 and BG505 at 25°C (Fig. 2B), the curve displays a downward trend at scattering vector values < 0.03Å (Fig. 2D). We performed a singular value decomposition of the static temperature series data set to examine the component curves that define the signal. One of the first two left vectors matches the difference curve between BG505 and CH505 consistent with the presence of differing open state propensity. The presence of the second curve suggests an Env shape difference occurring at elevated temperatures that is not different between these two Env isolate at ambient temperature. Together, these results demonstrate that the conformational transitions in CH505 SOSIP are sensitive to temperature and can be measured using SAXS.

### Time-resolved, temperature-jump SAXS reveals time-dependent changes in CH505 SOSIP SAXS Profiles

We next examined the CH505 SOSIP response to rapid system heating using TR, T-jump SAXS. We first optimized the initial, equilibrium temperature of our system. Based on our previous temperature-dependent binding experiments, we initiated temperature jumps from 40°C, 42°C, 44°C, and 46°C with a probe delay of 10 µs and 1 ms, and found that the difference signal in the region of interest (0.02-0.10 Å^-1^) was maximal at 44°C. We then determined the extent of laser-induced heating of the system when jumping from 42°C. Linear regression analysis (Supp. Fig. 4B) of the static SAXS difference profiles water ring maxima (1.5-2.5 Å^-1^) at Δ2°C, Δ5°C, Δ6°C, Δ8°C, and Δ11°C (Supp. Fig. 4A) indicated our system temperature jump was ∼Δ6°C leading to a final perturbed system temperature of ∼50°C. Next, to probe the CH505 SOSIP conformation at different times post-heating, we measured scattering at delay times of 500 ns, 1.5 µs, 3 µs, 5 µs, 10 µs, 50 µs, 100 µs, 500 µs, 1 ms, 10 ms, and 100 ms (Fig. 3A, Supp. Fig. 3B, E). The scattering difference curve measured at a post-laser temperature jump time of 500 ns shows a prominent negative peak between 0.02 and 0.05 Å^-1^ (Supp. Fig. 3E). This is suggestive of a process that occurs faster than our measurement dead time. This feature becomes increasingly prominent at greater time delays up to 1 ms with a marked increase between 0.02 and 0.05 Å^-1^ between 1.5 µs and 3 µs (Fig. 3A). Scattering at these low angles reports on the largest scale changes in the system (*122*) suggestive of a particle whose radius is increasing. A second signal at q = 0.07Å^-1^ becomes more prominent as the delay time is increased (Fig. 3A). Both the lowest scattering angle and 0.07 Å^-1^ features are consistent with the static temperature difference SAXS curves (Fig. 2B, D). At longer time delays of 10 ms and 100 ms, a period over which the system temperature is dropping (Supp. Fig. 4B), these difference features begin to shift toward zero (Fig. 3A), indicating the Env relaxes back to its initial state. Together, these results demonstrate the HIV-1 Env conformation exhibits reversible transitions on the microsecond timescale.

**Figure 3:**
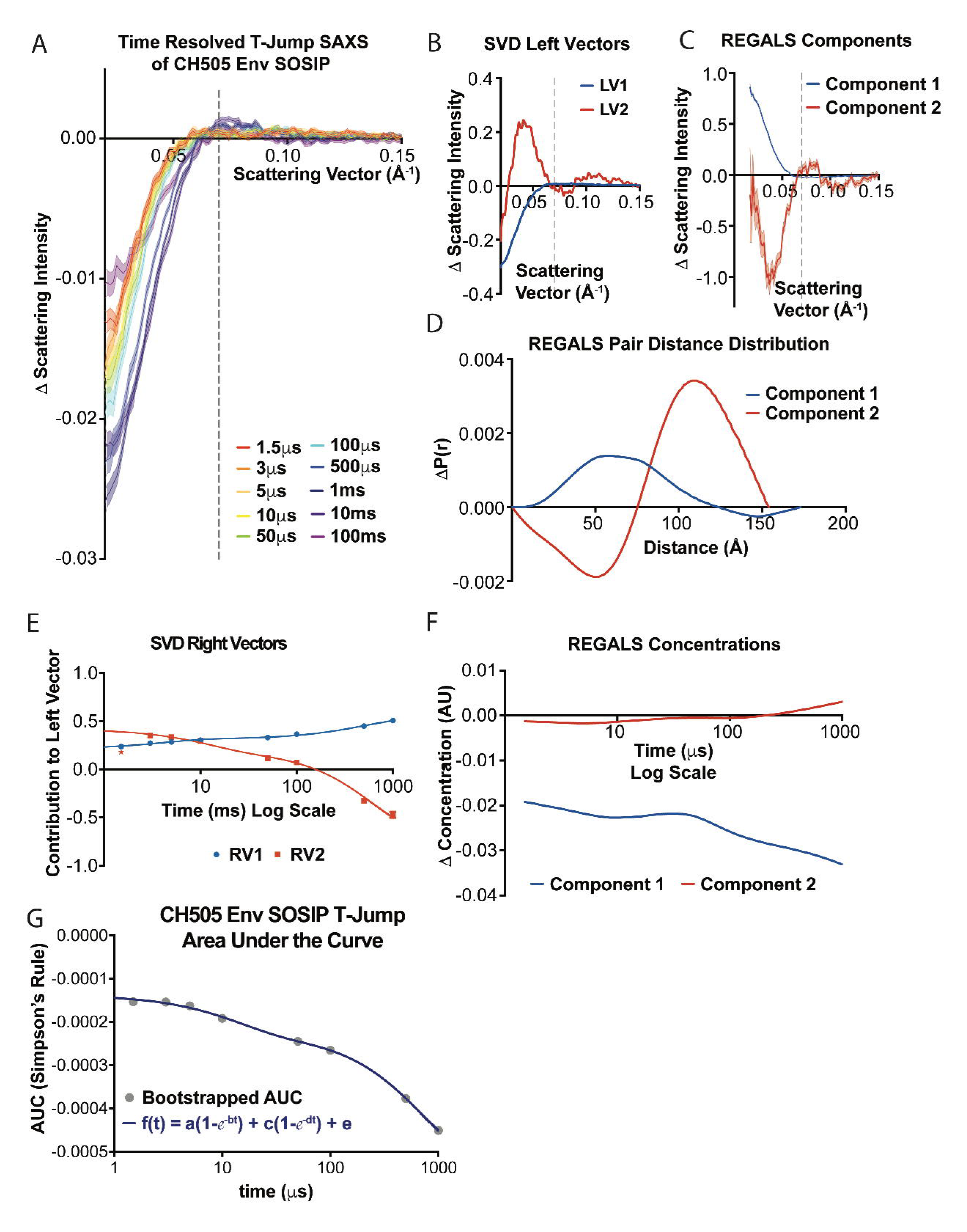
Time resolved, Temperature-Jump SAXS of HIV-1 Env Reveals Two Opening Transitions. **(A)** TR, T-Jump SAXS scattering difference curves for 1.5 µs (red) 3 µs (orange), 5 µs (light orange), 10 µs (yellow), 50 µs (green), 100 µs (cyan), 500 µs (blue), 1 ms (indigo), 10 ms (violet), and 100 ms (magenta) time delays. Dashed grey line at q=0.07 Å^-1^ indicates scattering difference feature peak. The shaded regions indicate the standard error of the arithmetic mean for the number of replicates indicated in the Methods section. **(B)** Singular value decomposition left vectors 1 (LV1, blue) and 2 (LV2, red). Dashed grey line at q=0.07Å^-1^ indicates the location of the feature peak. **(C)** Deconvoluted TR, T-Jump SAXS components 1 (blue) and component 2 (red) from REGALS decomposition SAXS difference curves in panel B. Dashed grey line at q=0.07 Å^-1^ indicates the location of the feature peak. **(D)** REGALS pair distance distribution for component 1 (blue) and component 2 (red). **(E)** The SVD right vectors 1 (RV1, blue) and 2 (RV2, red) showing the contribution of LV1 and LV2 at each time delay. The point shown as a red star was left out of the SVD fit to RV2. The arithmetic mean ± standard error of the mean (SEM) determined from bootstrapping is plotted. The error bars are smaller than the data point for most time delays. **(F)** The predicted concentrations of REGALS component 1 (blue) and component 2 (red). **(G)** The area under the curve (AUC) calculated according to Simpsons Rule for the TR, T-Jump SAXS difference curves shown in panel A and fit to a double exponential decay function. The arithmetic mean ± standard error of the mean (SEM) determined from bootstrapping is plotted. The error bars are smaller than the data point for most time delays.

### Env transitions through a rapidly forming intermediate

We next determined the number of distinct states observed and their exchange kinetics. For this, we first extracted component curves from the set of difference curves using singular value decomposition (SVD). This method splits the data into matrices of component curves, or left vectors, singular values, and contributions, or right vectors, that when multiplied, return the original curve (*123*). This acts to remove noise from the data and to separate signals into distinct processes. Here, we focused on difference curves between 1.5 µs and 1ms as these report primarily on structural transitions before cooling of the sample. The results indicated that there are two primary components in the time-resolved SAXS difference profiles (Supp. 5A). The first left vector is comprised of a negative peak between 0.02 and 0.05 Å^-1^ with a minor positive peak at 0.07 Å^-1^ (Fig. 3B). The second left vector shows features matching an inversion of the BG505 vs. CH505 difference curve (Fig. 3B). The right vectors report on the relative contributions of each component to each scattering curve and show that the first component transitions at both shorter and longer time delays while the second component transitions only at longer time delays (Fig. 3E, Table 2). Both the left vectors (Supp. Fig. 5B) and right vectors (Supp. Fig. 5C) for SVD components 3-8 fluctuate randomly about zero, indicating that these components do not contribute significantly to the time-resolved SAXS signal. We next fit kinetic models to each of the first two distinct component right vectors. A double exponential fit to the first component yields time constancts of 3.98 ± 0.07 µs and 507 ± 6 µs (Table 2). A double exponential fit to the second right vector yielded time constants of 12 ± 5 µs of 606 ± 145 µs (Table 2). The second component rates are consistent with the processes in the first component. We next determined time constants times from a double exponential fit to the area under each TR, T-Jump difference profile between 0.02 and 0.1 Å^-1^ (Fig. 3G). This yielded a time constant of 13.34 ± 0.04 µs and the other with a time constant of 708 ± 2 µs both of which are consistent with the values from the SVD fitting (Table 2). These models indicate that there is a rapid transition to an intermediate state followed by a second, slower transition.

Singular value decomposition does not necessarily decompose data into physically meaningful components (*124*). A recently developed algorithm, termed REGALS, deconvolves SAXS datasets by applying experimentally determined restraints to SVD deconvolutions so that the components are physically realistic (*124*). We performed a REGALS deconvolution on TR, T-Jump SAXS difference profiles to identify individual component scattering profiles in the CH505 SOSIP time-resolved SAXS data (Fig. 3C). We included two components as identified in both the SVD and area under the curve analyses with maximum dimensions of 173 Å and 154 Å, respectively, determined by the dimensions of Env structure models (Table3; Supp. Fig. 6A) and iterative model optimization. Fitting of the data yielded an overall *χ*^2^ of ∼1.0 (Supp. Fig. 5D). Each of the REGALS components (Fig. 3C) displayed similarities to their respective SVD components (Fig. 3B) despite being reflected about the X-axis. In addition to splitting the scattering difference curves into distinct components, REGALS returns pair distance distributions describing component particle atom-atom distance differences (Fig. 3D).

The distance distribution difference for component 2 indicated structural changes occur on a relatively large range up to ∼120 Å, with marked reductions in pairwise distances between 25 Å and 75 Å and concomitant increases between 80 Å and 120 Å. The largest distance changes for component 1 were over a smaller range with a mean of ∼55 Å (Fig. 3D). The concentration of component 2 begins to increase at longer time delays while component 1 decreases at early and long time delays (Fig. 3F), a feature consistent with the changing contributions of the components determined by the SVD right vector analysis (Fig. 3E). These results are consistent with a model of an early intermediate structural state giving way to a more structural state in which large scale domain movements occur after a delay of nearly one millisecond.

### Trimer apex contacts break before gp120 rotation

We next asked how the transition and states observed here relate to known HIV-1 Env ectodomain structures and dynamics. Previous smFRET studies have shown that an asymmetric intermediate connects the closed and open configurations (*69-72*). Transitions from the ground state to this asymmetric state and from this state to the open state often occur within the dead time of the measurement (*72*) which is consistent with the processes measured here (Fig. 3G, Table 2). The asymmetric state is characterized by a single protomer open state with two protomers in a distinct conformation. Based on the correlation between open-state targeting antibody binding and static SAXS at increasing temperature, the first transition observed in the TR, T-jump SAXS dataset corresponds to transitions that are initiated from a closed state. Two distinct structural states of SOSIP Envs have been observed previously including open-occluded states in which each gp120 domain is rotated outward from the primary trimer axis without major rearrangements in V1/V2 or V3, and an open state in which the gp120s domains are rotated outward and V1/V2 and V3 have rearranged (Fig. 1B). We first asked whether our measures could effectively distinguish between theoretical SAXS difference curves calculated between a closed state model and either an open-occluded model or an open model (Fig. 4A-C; Supp. Fig. 6A). An SVD analysis of the closed, open-occluded, and open model theoretical SAXS curves (Supp. Fig. 6B) indicated two primary signals consistent with the components extracted from the TR, T-Jump experimental difference curves (Supp. Fig. 6C). The SAXS difference curve for a transition from closed to open occluded resembles the SVD and REGALS components 2, with a prominent negative and positive peak at low scattering angle (Fig. 4A). The closed to open SAXS difference curve was dominated by a single positive peak (Fig. 4B). The P(r) function difference curves for each were like the REGALS component P(r) difference curves, albiet inverted across the distance axis (Fig. 4A and B). The open-occluded vs. open SAXS and P(r) function difference curves did not show similarities to any of the SVD or REGALS curves (Fig. 4C). This indicates transitions between these two states did not occur or were not resolved in the dataset. Together, these modeling results suggest the second, slower transition involves gp120 rotation.

**Figure 4:**
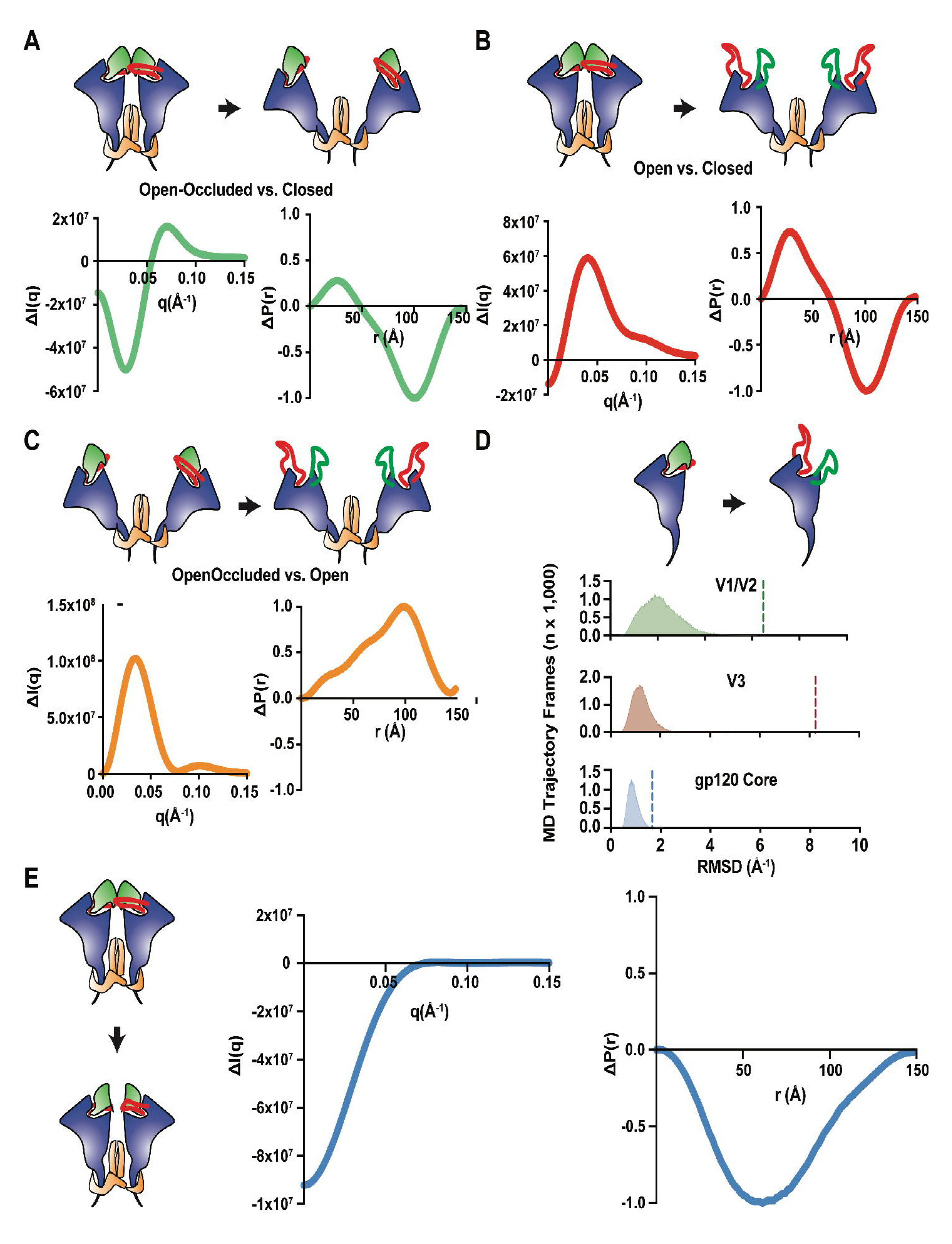
The fast intermediate corresponds to a loss of Env trimer apex contacts. **(A)** A cartoon diagram representing the closed to open-occluded transition using the same coloring as in Figure 1 (top). The theoretical SAXS difference curve (*bottom, left*) and the theoretical pair distance distribution difference curve (*bottom, right*) for the closed to open-occluded transition. **(B)** A cartoon diagram representing the closed to open transition using the same coloring as in Figure 1 (*top*). Theoretical scattering difference curves (*bottom, left*) and pair distance distribution difference curve (*bottom, right*) for the closed to open conformational transition. **(C)** A cartoon diagram of the Env SOSIP transition from the open-occluded conformation to the open conformation (*top*). The theoretical SAXS scattering difference curve (*bottom, left*) and pair distance distribution difference curve (*bottom, left*) for the open-occluded to open transition. **(D)** The carton diagram depicts the V1/V2 and V3 rearrangement in a gp120 monomer (*top*). The RMSD (*bottom*) for V1/V2 (green), V3 (blue), and gp120 core (blue) b-sheet a-carbons for an aggregated 250 5 µs MD simulations. The dashed lines represent the RMSD for the given domain between the starting and end conformations depicted in the diagram. **(E)** A cartoon diagram depicting the release of the Env SOSIP trimer apex contacts (*left*). The theoretical SAXS scattering difference curve (*middle*) and the pair distance distribution difference curve (*right*).

We next examined the structural transition involved in the rapidly forming intermediate. The first REGALS component P(r) difference function involves a substantial loss of scattering centers within a range of 0-100 Å but without concomitant increases in more distant scattering centers. This is consistent with a loss of scattering density in the Env. The HIV-1 Env is a heavily glycosylated protein with a dense network of conformationally dynamic interactions (*125-127*). We, therefore, asked whether changes in the glycan shield could explain the two SVD and REGALS components. An SVD analysis based on theoretical SAXS profiles for non-glycosylated and Man9 glycosylated Env closed, open-occluded, and open models indicated the overall features between the two are similar with shifts in the scattering vector position of the peaks that are related to differences in particle size between the two sets (Supp. Fig. 6E). Importantly, the non-glycosylated models display the SAXS features of interest, suggesting that the glycans alone do not give rise to the observed differences signal.

We next asked whether the intermediate state involves changes in the apex conformation. Isolated gp120 domains in solution are known to expose the bridging sheet epitope targeted by the CCR5 binding site mAb, 17b, as well as the V3 loop targeting antibodies such as mAbs, 19b, F39F, and 3074 (*128*). In the absence of Env trimer apex contacts, a free gp120 domain with the V1/V2 and V3 regions in their observed closed state structural arrangement should therefore spontaneously sample an open-like gp120 state in which the bridging sheet is formed. We collected two-hundred and fifty independent 5 µs molecular dynamics simulation trajectories run at 50°C to examine the closed-state fold stability at a timescale similar to that of the observed intermediate. The root mean square deviation (RMSD) of a-carbon atoms in the gp120 core b-sheets (0.9 ± 0.2 Å) was markedly smaller than both V3 (1.3 ± 0.4 Å) and V1/V2 (2.1 ± 0.8 Å) (Fig. 4D). These differences occurred due to an order-to-disorder transition in the V1/V2 and V3 loops that form apex loop contacts in the closed state trimer. This led us to ask whether an order-to-disorder transition in the apex could explain the change in scattering observed in the fast-forming intermediate. The loss of scattering density suggested by the REGALS component 1 P(r) difference function is, indeed, consistent with an order-to-disorder transition. This is due to a reduced density differential for a packed protein fold and a well-hydrated, disordered chain. We, therefore, asked whether a structure missing the gp120-to-gp120 contacting loops could give rise to the observed scattering difference curves and ΔP(r) function. The difference profile for both the scattering and P(r) functions calculated for models missing residues 160-171 and 306-317 indeed matched the observed REGALs component 1 (Fig. 4E). This is reflected as a reduction in scattering density and likely corresponds to a transition to a disordered state for this portion of the trimer. Together, these results are consistent with a model of a T-jump transition that first eliminates contacts at the trimer apex that is followed later by gp120 rotation.

### Interprotomer Disulfide Bonds Stabilize the Closed Env Trimer

The TR, T-Jump SAXS results indicate an intermediate structural state precedes movements to a more open configuration. A recent macaque immunization study found that, despite immunogen stabilization in a prefusion closed conformation, antibodies targeting open-occluded states were induced (*43*). Our modeling of the SAXS data suggests changes in gp120- to-gp120 apex contacts occur first and at a rate that is inconsistent with major rearrangements in the allosteric machinery including intra-protomer V1/V2 and V3 displacement. Thus, we argue that blocking access to the fast-forming intermediate would prevent the Env from transitioning to the open occluded conformation. We, therefore, used a combination of *in silico* tools to identify possible inter-protomer apex stabilizing mutations. From this analysis, we identified a V127C-D167C disulfide that stapled V1/V2 contact regions between gp120 protomers at V1/V2 contact regions in a CH505 SOSIP trimer design parent (*42*). This design was further stabilized using the previously reported F14 (*49*) and SOSIP 2P (*129*) mutations to improve expression and folding in addition to a CD4bs antibody binding enhancing N197D glycan deletion mutation (*117*). Even though a closed trimers Env conformation was identified by NSEM, SDS-PAGE analysis as well as binding to 19b indicated that the Env SOSIP preparation included a population with improperly formed inter-protomer disulfides and was exposing the epitope for 19b (Supp. Fig. 7). Single-particle cryo-electron microscopy (cryo-EM) analysis a CH505 apex stapled SOSIP design bound to CH235.12 yielded two distinct populations, including a closed state and another state lacking well-defined density in the V1/V2 apical region (Fig. 5C, Supp. Fig. 8B). To eliminate the open state, we further purified the Env SOSIP sample by negative selection using the V3 loop targeting 3074 antibody (*130*). This resulted in the elimination of the non-inter-protomer linked bands in SDS-PAGE, reduction of 19b binding, and enhancement of trimer specific, gp120/gp41 interface targeting PGT151 binding (Fig. 5B, Supp. Fig. 7A). The apex stapled design displayed a marked increase in thermal denaturation inflection temperature (T_i_ +7.6°C; Fig. 5A, Supp. Fig. 7D). The closed state configuration of the inter-protomer disulfide stapled design displayed lower resolution density at the inter-protomer apex loop contacts suggestive of conformational variability at this site (Supp. Fig. 8F). Density consistent with the formation of a disulfide between the protomers was nevertheless visible at a lowered map contour level (Supp. Fig. 11A). Together, these results show that disulfide linkages between protomers at the trimer apex can effectively stabilize the trimer.

**Figure 5:**
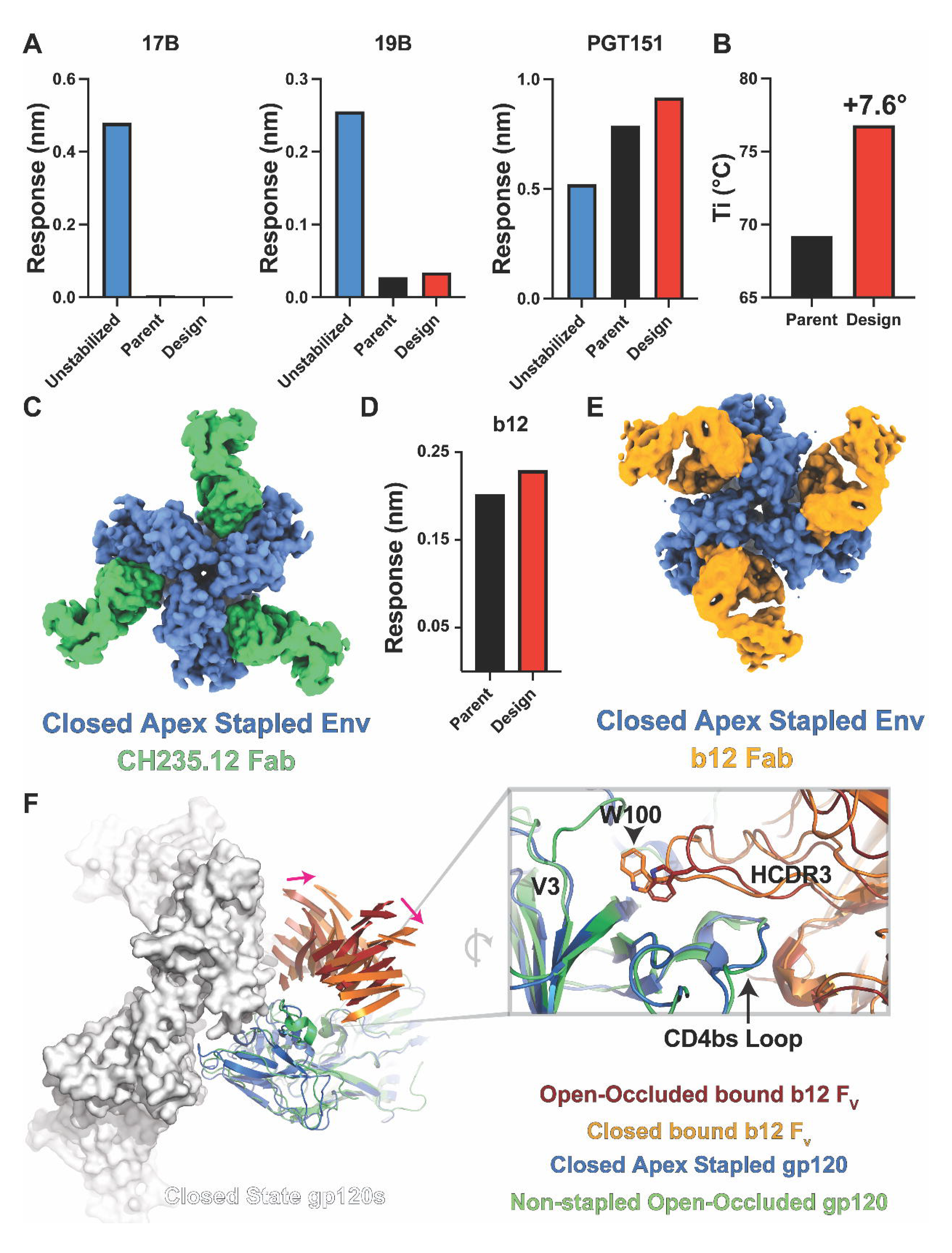
Interprotomer Disulfide Bonds Stabilize the Closed Env Trimer. **(A)** Differential fluorescence thermal denaturation inflection point temperature for the stabilized parent CH505 SOSIP and the inter-protomer disulfide stapled CH505 SOSIP. **(B)** Binding responses for an unstabilized CH505 SOSIP, the stabilized CH505 parent SOSIP, and the inter-protomer disulfide stapled CH505 SOSIP design interacting with the co-receptor binding 17b, V3-loop binding 19b, and the closed state apex interactive, trimer specific PGT145 MAbs. **(C)** A gaussian filtered map of the inter-protomer disulfide stapled CH505 SOSIP design bound to the CH235.12 Fab. **(D)** Binding responses for PGT151 captured stabilized CH505 parent SOSIP and inter-protomer disulfide stapled CH505 SOSIP design interacting with the b12 Fab. **(E)** A gaussian filtered map of the inter-protomer disulfide stapled CH505 SOSIP design bound to the b12 Fab. **(F)** (left) Structure comparison between the closed state b12 bound inter-protomer stapled CH505 SOSIP design and the open-occluded state b12 bound B41 isolate SOSIP. A single b12 bound gp120 domain from B41 is aligned to a closed state CH505 design gp120 to highlight the shift in the b12 Fv position (only β-sheets shown for clarity; pink arrows indication difference direction). Two additional gp120 domains shown as surface to highlight potential for b12 clashes. (right) Alignment of the b12 bound apex stapled CH505 SOSIP design and B41 isolate gp120 domains highlighting differences in the HCDR3 position.

We next asked whether open-occluded state antibody binding was eliminated in the inter-protomer disulfide-linked trimers. A previous structural study showed that the CD4bs b12 antibody interacted with the B41 isolate Env ectodomain in an open-occluded conformation (*10*). Another study showed that the capture of a BG505 isolate Env ectodomain with PGT145 eliminated b12 binding, presumably through locking the trimer apex in the close configuration (*43*). Interestingly, the apex stapled CH505 SOSIP design binding with the b12 antibody Fab by BLI was like that of the non-apex stapled construct (Fig. 5D). Examination of the b12 bound structures of each by NSEM indicated that in the non-apex stapled construct apex contacts were broken, consistent with an open-occluded state, while the apex contacts in the apex stapled construct remained intact (Supp. Fig. 9). Model fitting of b12 into the apex stapled density suggested that a substantial shift in its angle of approach permitted interaction of b12 with this apex-stapled construct fixed in the closed configuration and unable to access the open-occluded conformation preferred by b12 (Supp. Fig. 9). We obtained a 4.5 Å resolution cryo-EM reconstruction of the b12 bound apex stapled construct to examine these changes in greater detail (Supp. Fig. 10). The map showed that the prefusion, closed configuration is retained with b12 interaction and that the b12 angle of approach is indeed substantially shifted compared to a previously determined open-occluded state b12 bound B41 isolate Env SOSIP (Fig. 5E, F) (*10*). This results in a shift of the HCDR3 loop bringing W100 nearer the V3 loop (Fig. 5F). The overall fold of the gp120 domain is similar, with a gp120 RMSD of ∼1.3 Å (Supp. Fig. 11B). The shifted position appears to be facilitated through rearrangements in the β20-β21 and b12 contact with V1/V2 region relative to the B41 isolate bound, open-occluded state structure (Supp. Fig. 11C). These results show that disulfide linkages at the trimer apex between protomers can effectively eliminate movements to an open-occluded, intermediate state.

## Discussion

Here, we show that the CH505 Env SOSIP conformation transitions to an open-occluded-like state through an intermediate that forms on the tens of microseconds timescale. The gp120 MD simulations show that the closed state fold from which they were initiated is stable on the timescale of the fast-forming intermediate apart from the interprotomer contacting gp120 loops. These loops showed an order-to-disorder transition in the simulations. This led us to test a model for early intermediate formation involving loss of scattering density at the apex loop contacts. We found that this model is consistent with both our SVD and REGALS analysis suggesting these loops indeed undergo an ordered to disordered transition. The second, slower transition is consistent with scattering profile differences between the CH505 and BG505 SOSIP Envs and open-state antibody binding. Thus, this state likely corresponds with an Env in which gp120 is rotated outwards, giving rise to a greater distance between the gp120 apex residues. It is not clear from the scattering data here, however, to what extent the V1/V2 and V3 rearrangements have occurred or the extent of gp120 rotation.

Most stabilizing designs focus on disabling allosteric rearrangements coupled with V1/V2 rearrangement and improving gp120 to gp41 interactions (7, 18, 20, 25, 49, 53, 54, 63, 73-79). The induction of antibodies that target the open-occluded state of Env by vaccination with pre-fusion stabilized trimers and the presence of an intermediate consistent with this state here suggests stabilization between apex contacts is needed to eliminate shifts in gp120 domain position. As blocking transitions out of the closed state is essential for inducing neutralizing antibodies by vaccination, we used this information to design an Env SOSIP with a disulfide staple at the apex to eliminate this disorder transition. This effect was confirmed by the closed state-bound configuration of the b12-neutralizing antibody. Together, these results show that microsecond dynamics control movement between the closed and open states.

When interpreting low-resolution scattering results, it is important to consider alternative explanations of the data. The Env glycan shield likely plays a major role in determining the density of this solvent shell and, in combination with the possibility of differential glycan interactions, complicates differentiating transitions that involve protein and those that may involve only glycan. We find that modeling the trimer in different combinations of closed and open states here with and without glycans yields nearly identical difference curve features. This suggests the early transitions we observe are indeed protein related. The solvent shell of non-glycosylated proteins responds to the temperature jump within ∼250 ns (*131*). The relaxation timescale for individual glycan motions is typically under one microsecond at a given temperature (*132*) which together suggests that any changes in the glycan in response to the temperature jump would likely occur well below our observed intermediate timescale. These considerations together lead us to conclude that a substantial portion of time-dependent scattering differentials we observe are associated with protein movements.

It is also important to consider that, though the Env SOSIP used here was extensively purified and characterized, we cannot entirely rule out the possibility that multiple, yet unidentified, folded species exist in our samples. This too could lead to two different transition rates that, rather than sequential movements, are related to relative transition propensities between two different folded states. In the case of our apex stapled design, a differing folded state population was readily identified in our cryo-EM 3D classification. No such state has been identified in any of our related SOSIP constructs (*42, 49, 90, 133*), nor were substantial lot-to-lot quality control variations observed that would indicate fold instability. Further, a different fold would likely result in a distinct thermal denaturation profile, so a discernable bi or multimodal denaturation profile would be expected. This was not observed for our samples here. We, therefore, conclude that the transitions we observe are indeed related to sequential events of a single-fold species.

The findings from this study show that the transition from a closed state to one in which the gp120 domains rotate outward, away from the timer central axis involves sequential movements. Rather than gp120 rotation occurring simultaneously with apex contact breakage, the gp120 apex loop contacts first become disordered. Our results suggest this occurs without complete V1/V2 rearrangements that result in the formation of the bridging sheet. This is consistent with previous studies showing V3 loop exposure can occur absent exposure of the bridging sheet (*57*). The transition to a more open configuration occurred at nearly two orders of magnitude longer timescale. With contacts at the apex broken, control of this transition must involve the remaining gp120 to gp41 contacts. At equilibrium, this sequential ordering of each transition creates a kinetic switch where gp120 rotation is controlled by the joint probability of both the order-to-disorder apex transition and breakage of the remaining gp120 to gp41 contacts occurring. Changes in the relative probabilities of these transitions likely occur between different isolates and may correlate with sensitivities to receptor-induced conformational transitions.

The relationship between the transitions we observe here and the native, full-length gp160 HIV-1 Env is an open question. Native Env trimers are membrane-embedded, which may alter the conformational transitions between Env structural states and their kinetics relative to the transitions measured here in the soluble gp140 construct. Membrane-embedded gp160 Env is known to transition between multiple structural states similar to the soluble SOSIP gp140 constructs (*2, 69, 70, 134*) and also shares structural similarity with gp140 SOSIP structures (*46, 56*). Thus, the results presented here are likely relevant for gp160 Env. The specific relationship between previously observed states is nevertheless unknown. These are important questions as any susceptibility to movement beyond a prefusion closed state could be exploited by maturing antibodies in the context of vaccination. These states are rare in tier 2 viruses absent the CD4 receptor. The impacts on vaccination outcome are two-fold, poorly neutralizing antibodies targeting open-occluded states are more likely to be induced and antibodies with broad neutralizing potential can acquire mutations favoring these states rather than neutralization critical mutations, thus limiting their development toward breadth. The apex stapled design presented here prevents the movement of gp120, thus eliminating these possibilities. Consistent with impacts on antibody interactions, the NSEM and cryo-EM structures of b12 revealed marked changes in the antibody angle of approach with the apex staple compared to a non-stapled construct.

In summary, these results show that transient intermediate states observable only on a microsecond scale play an essential role in controlling HIV-1 Env conformation. Blocking these early transitions is likely an important consideration in vaccine development efforts to ensure maturing antibodies remain on track to develop neutralization breadth.

## Acknowledgments

This project was supported by NIH, NIAID, Division of AIDS Consortia for HIV/AIDS Vaccine Development (CHAVD) Grant UM1AI144371 (B.F.H), R01AI145687 (P.A.), U54AI170752 (R.C.H. and P.A.), and Translating Duke Health Initiative (R.C.H. and P.A.). A portion of this work was conducted at the Advanced Light Source (ALS), a national user facility operated by Lawrence Berkeley National Laboratory on behalf of the Department of Energy, Office of Basic Energy Sciences, through the Integrated Diffraction Analysis Technologies (IDAT) program, supported by DOE Office of Biological and Environmental Research. Additional support comes from the National Institute of Health project ALS-ENABLE (P30 GM124169) and a High-End Instrumentation Grant S10OD018483. This research also used resources of the Advanced Photon Source, a U.S. Department of Energy (DOE) Office of Science User Facility operated for the DOE Office of Science by Argonne National Laboratory under Contract No. DE-AC02- 06CH11357. The use of BioCARS was also supported by the **<u>National Institute of General Medical Sciences</u>** of the **<u>National Institutes of Health</u>** under grant number P41 GM118217. The content is solely the responsibility of the authors and does not necessarily represent the official views of the National Institutes of Health. Time-resolved set-up at Sector 14 was funded in part through a collaboration with Philip Anfinrud (NIH/NIDDK). Prof. Lin X. Chen of Northwestern University provided the heating cell for TR, T-Jump SAXS experiments.

## Author Contributions

A.L.B. and R.C.H. designed, conducted, and interpreted the TR, T-jump SAXS experiments. A.L.B. developed the SAXS analysis code and analyzed SAXS data. C.S., A.W., and Y.B. prepared all proteins for this study. C.S. and Y.B. conducted SAXS experiments. C.S. collected and analyzed BLI data. R.J.E. and K.M. collected and analyzed NSEM data. R.C.H. collected the cryo-EM data. R.C.H. and P.A and analyzed the cryo-EM data. R.C.H. designed the apex-stapled immunogen. All authors read and edited the manuscript.

## Competing Interest

The authors declare the following competing interests: A patent application covering HIV-1 Envelope modifications based on this study has been submitted by Duke University.

## Data and Code Availability

All data and code from this manuscript are available upon reasonable request.

