## Supplemental Figures and Tables for "Microsecond dynamics control the HIV-1 envelope conformation"

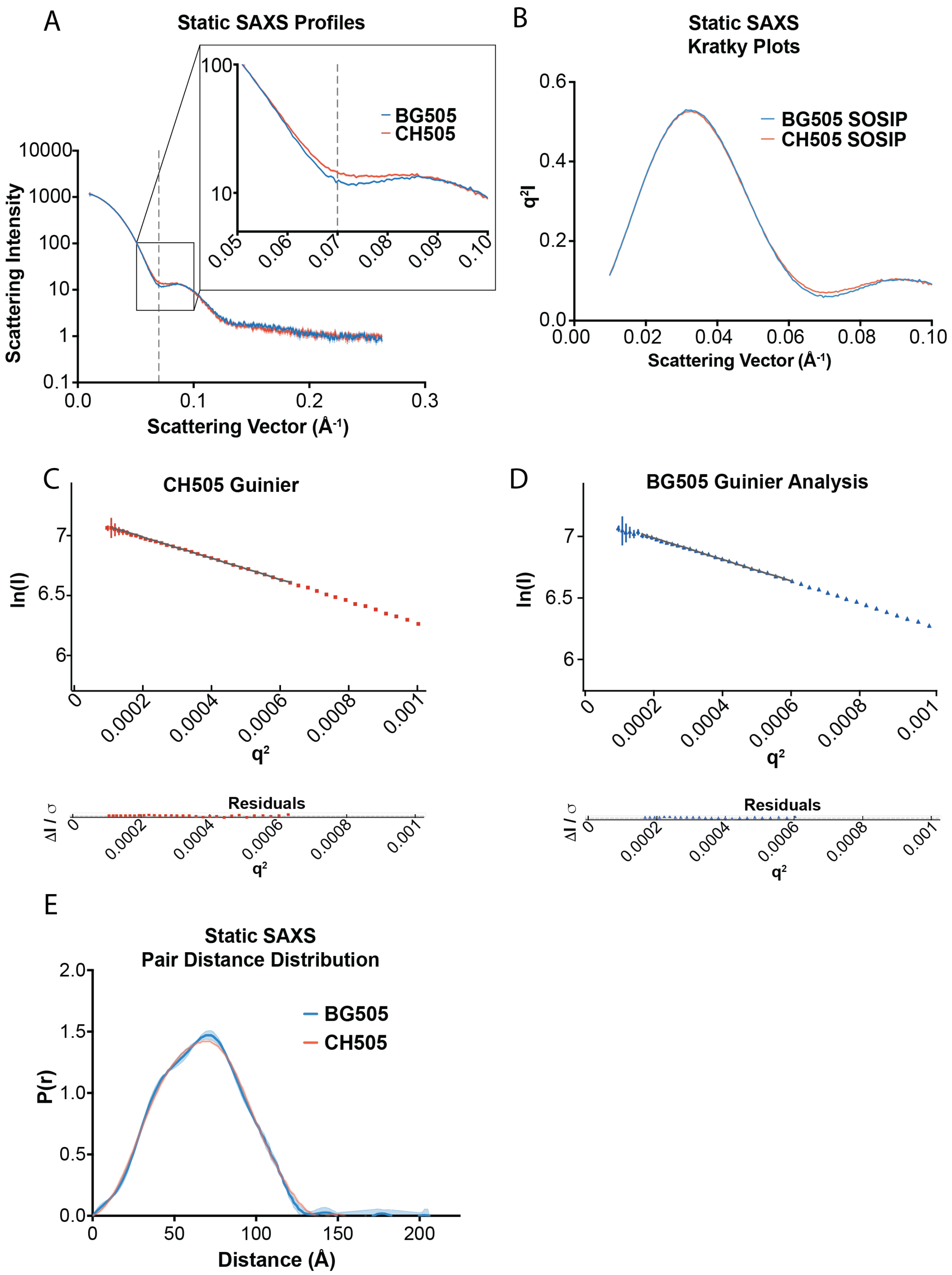
A
Static SAXS Profiles
Scattering Intensity
Scattering Vector (Å⁻¹)
BG505
CH505
B
Static SAXS
Kratky Plots
q²I
Scattering Vector (Å⁻¹)
BG505 SOSIP
CH505 SOSIP
C
CH505 Guinier
ln(I)
q²
Residuals
ΔI / σ
D
BG505 Guinier Analysis
E
Static SAXS
Pair Distance Distribution
P(r)
Distance (Å)

**Supplemental Figure 1: Different HIV-1 Env SOSIP isolates show differences in SAXS profiles. (A)** Static SAXS profiles for BG505 Env SOSIP (blue) and CH505TF Env SOSIP (red). The scattering intensity is expressed on the y-axis as a function of the scattering vector in $Å^{-1}$. The box shows the region of the scattering profile displayed in the inset. The grey dashed line indicates the feature peak at q = 0.07 $Å^{-1}$. **(B)** The Kratky plots for BG505 SOSIP (blue) and CH505TF SOSIP (red). Guinier analysis of CH505 Env **(C)** and BG505 Env SOSIP **(D)** SAXS profiles. **(E)** The pair distance distribution (P(r)) for BG505 Env SOSIP (blue) and CH505TF Env SOSIP (red).

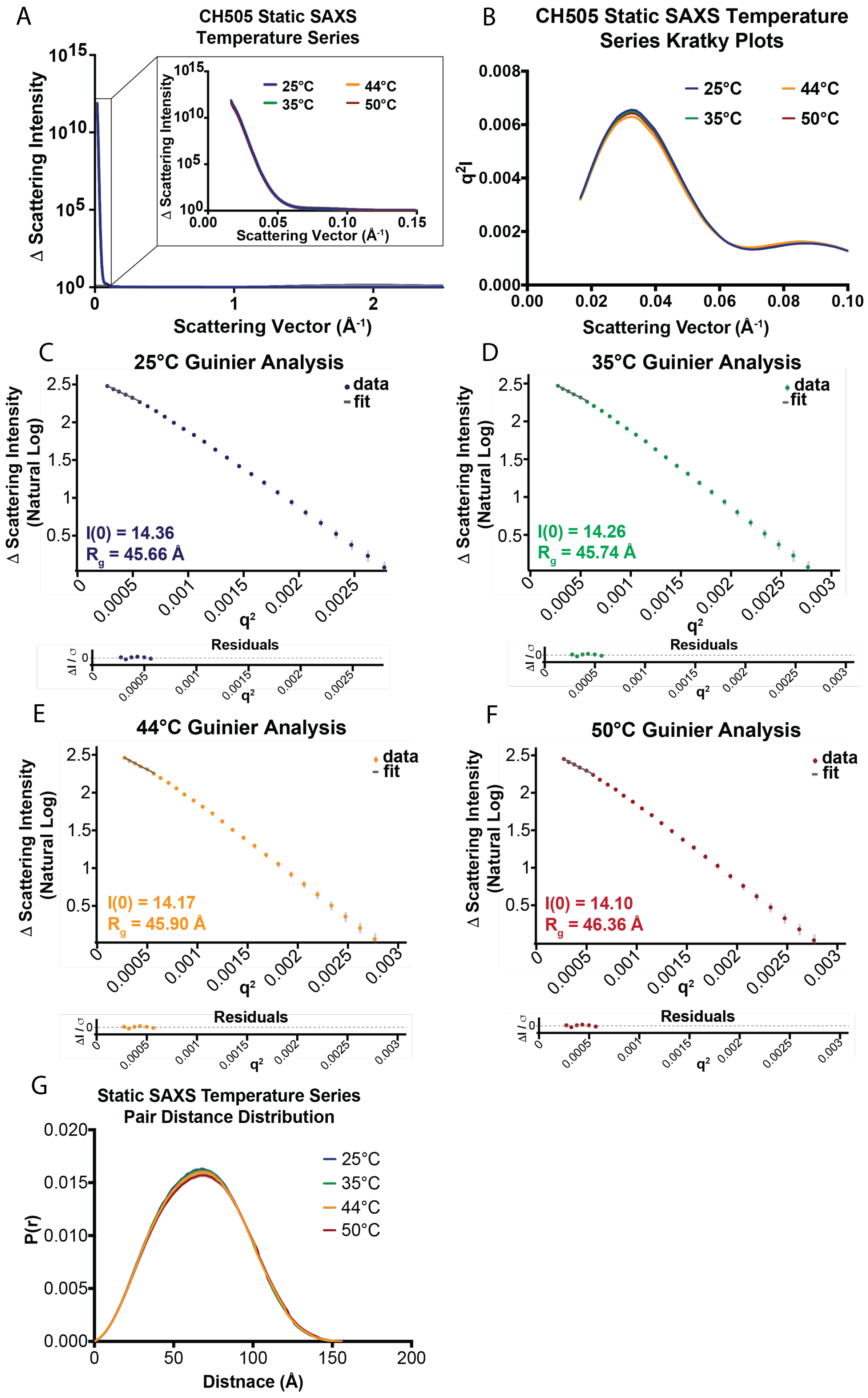
A
CH505 Static SAXS Temperature Series
Δ Scattering Intensity
Scattering Vector (Å-1)
25°C
35°C
44°C
50°C
B
CH505 Static SAXS Temperature Series Kratky Plots
q2I
Scattering Vector (Å-1)
C
25°C Guinier Analysis
I(0) = 14.36
Rg = 45.66 Å
D
35°C Guinier Analysis
I(0) = 14.26
Rg = 45.74 Å
E
44°C Guinier Analysis
I(0) = 14.17
Rg = 45.90 Å
F
50°C Guinier Analysis
I(0) = 14.10
Rg = 46.36 Å
Δ Scattering Intensity (Natural Log)
data
fit
q2
Residuals
ΔI / σ
G
Static SAXS Temperature Series Pair Distance Distribution
P(r)
Distnace (Å)

**Supplemental Figure 2: HIV-1 Env SOSIP is stable under SAXS at elevated temperatures.** **(A)** Static SAXS profiles for CH505TF Env SOSIP at 25°C (blue), 35°C (green), 44°C (yellow), and 50°C (red). The scattering intensity is expressed on the y-axis as a function of the scattering vector in $Å^{-1}$. The box shows the region of the scattering profile displayed in the inset. **(B)** The Kratky plots for CH505TF SOSIP at 25°C (blue), 35°C (green), 44°C (yellow), and 50°C (red). **(C)** Guinier analysis of CH505TF Env SOSIP SAXS profiles at 25°C. **(D)** The Guinier analysis of CH505TF Env SOSIP SAXS profiles at 35°C. **(E)** The Guinier analysis of CH505TF Env SOSIP SAXS profiles at 44°C. **(F)** The Guinier analysis of CH505TF Env SOSIP SAXS profiles at 50°C. **(G)** The pair distance distribution (P(r)) for CH505TF Env SOSIP at 25°C, 35°C, 44°C, and 50°C in blue, green, yellow, and red, respectively.

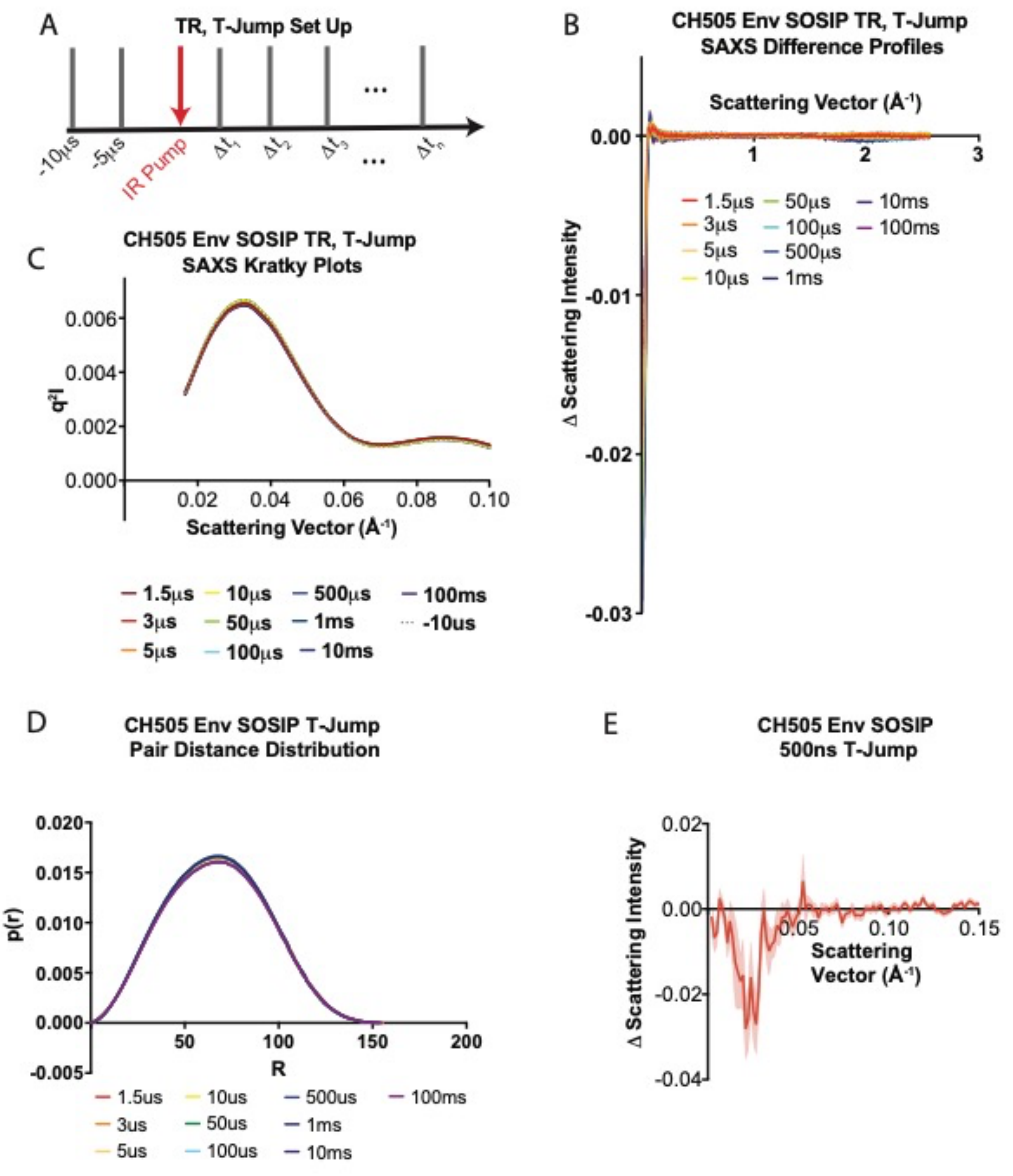

**Supplemental Figure 3: CH505TF Env SOSIP shows time-dependent changes in SAXS profiles. (A)** Experimental protocol for the pump-probe set up. The horizontal black arrow represents the progression of time during the experiment. The grey bars represent the X-ray probe step, and the red bars represent the infrared laser pump step. For the time resolved (TR) temperature-jump (T-jump) SAXS experiments, the X-ray scattering was collected at 2 time points (-10μs and -5μs) prior to the infrared laser pump and then the IR pump step is interleaved with the X-ray probe at various time delays after the IR pump. The time delays measured for this experiment range from 1.5 μs to 100 ms as well as 500 ns **(B)** TR, T-Jump SAXS scattering difference curves for 1.5 μs (red), 3 μs (orange), 5 μs (light orange), 10 μs (yellow), 50 μs (green), 100 μs (cyan), 500μs (blue), 1 ms (indigo), 10 ms (violet), and 100 ms (magenta) time delays. Dashed grey line at $q=0.07Å^{-1}$ indicates the location of the feature of interest and shaded regions represent the standard error of the arithmetic mean. **(C)** The Kratky plots for CH505TF Env SOSIP at 1.5 μs (red), 3 μs (orange), 5 μs (light orange), 10 μs (yellow), 50 μs (green), 100 μs (cyan), 500 μs (blue), 1 ms (indigo), 10 ms (violet), and 100 ms (magenta) time delays. The Guinier analysis of CH505TF Env SOSIP SAXS profiles at 44°C. **(F)** TR, T-Jump SAXS scattering difference curves for 500ns with the change in scattering intensity plotted as a function of the scattering vector in $Å^{-1}$. The shaded areas represent the standard error of the arithmetic mean.

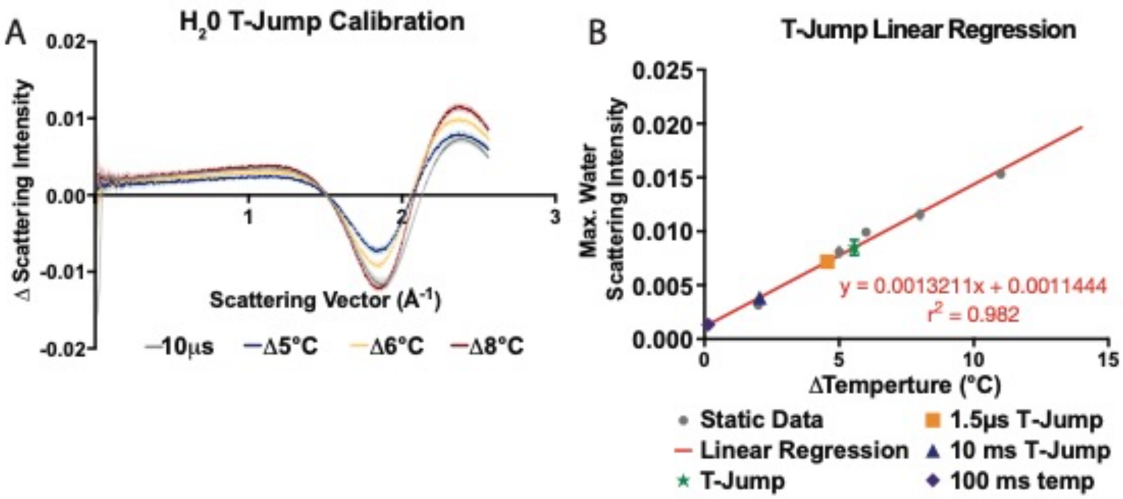

**Supplemental Figure 4: Calibration of TR, T-Jump. (A)** Static SAXS scattering difference curves for Δ5 (blue), Δ6 (yellow), Δ8 (red), and 10μs $H_2O$ T-Jump (grey). The change in scattering intensity is plotted as function of the scattering vector in $Å^{-1}$. **(B)** Linear fit of the difference curves in panel A. The maximum water scattering intensity is plotted as a function of the temperature difference. The grey circles represent the experimental data, and the red line is the linear regression fit to the experimental data. The fitted equation and the correlation coefficient are shown in red. Estimates for the 10μs $H_2O$ T-Jump, 1.5 μs CH505TF Env SOSIP T-Jump, 10 ms CH505TF Env SOSIP T-Jump, and the 100 ms CH505TF Env SOSIP T-Jump are shown by the green star, yellow square, blue triangle, and purple diamond, respectively.

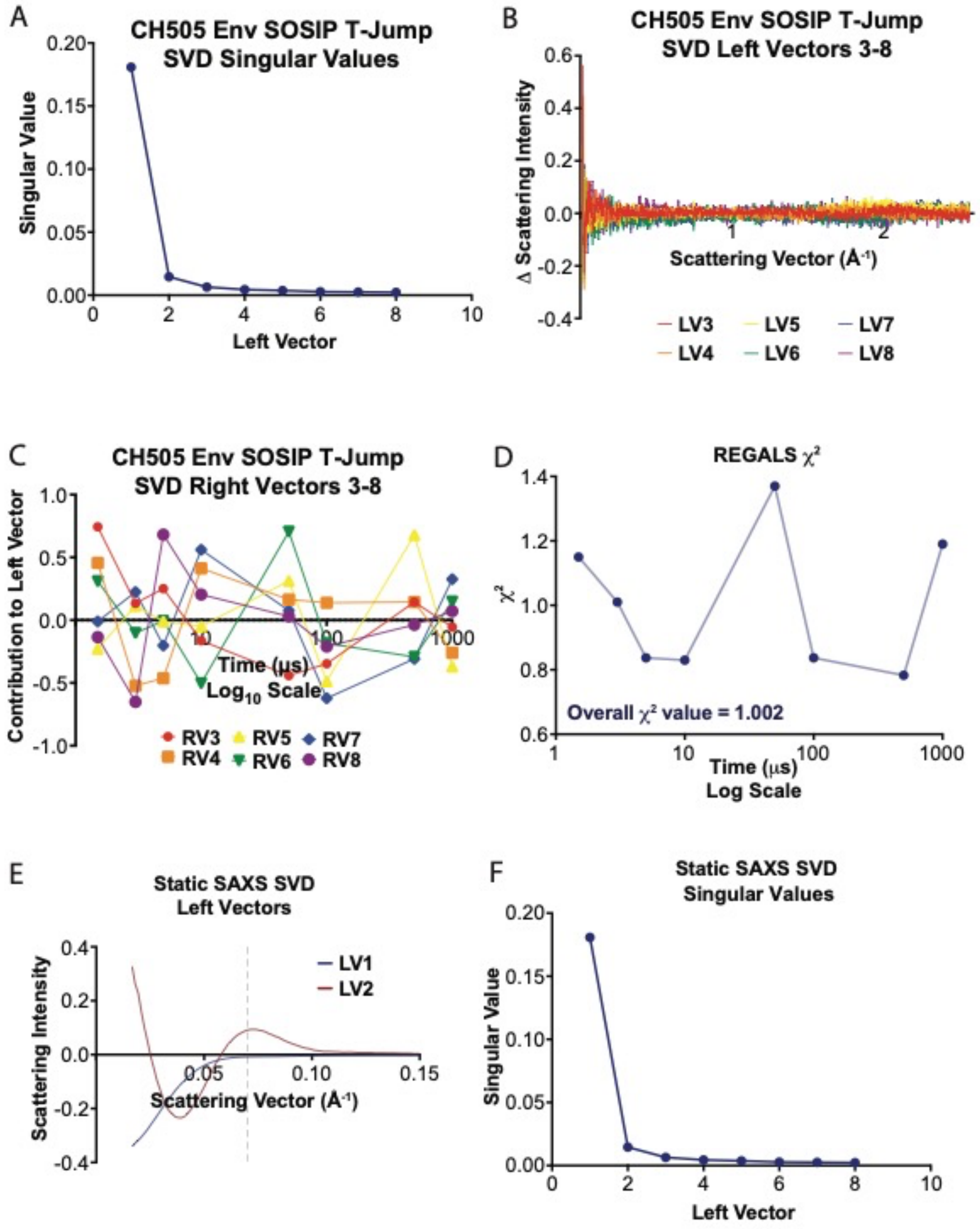

**Supplemental Figure 5: Decomposition of CH505TF Env SOSIP TR, T-Jump SAXS Difference Profiles. (A)** The singular values of singular value decomposition (SVD) on

CH505TF Env SOSIP TR, T-Jump difference profiles plotted as a function of the extracted SVD vector. **(B)** CH505TF Env SOSIP TR, T-Jump SVD left vectors 3-8. **(C)** CH505TF Env SOSIP TR, T-Jump SVD right vectors. **(D)** $\chi^2$ values of the REGALS fit to the TR, T-Jump SAXS data. **(E)** The SVD left vectors for CH505 Env SOSIP static SAXS temperature series. **(F)** The SVD singular values for the SVD components extracted from the CH505 Env SOSIP static SAXS temperature series.

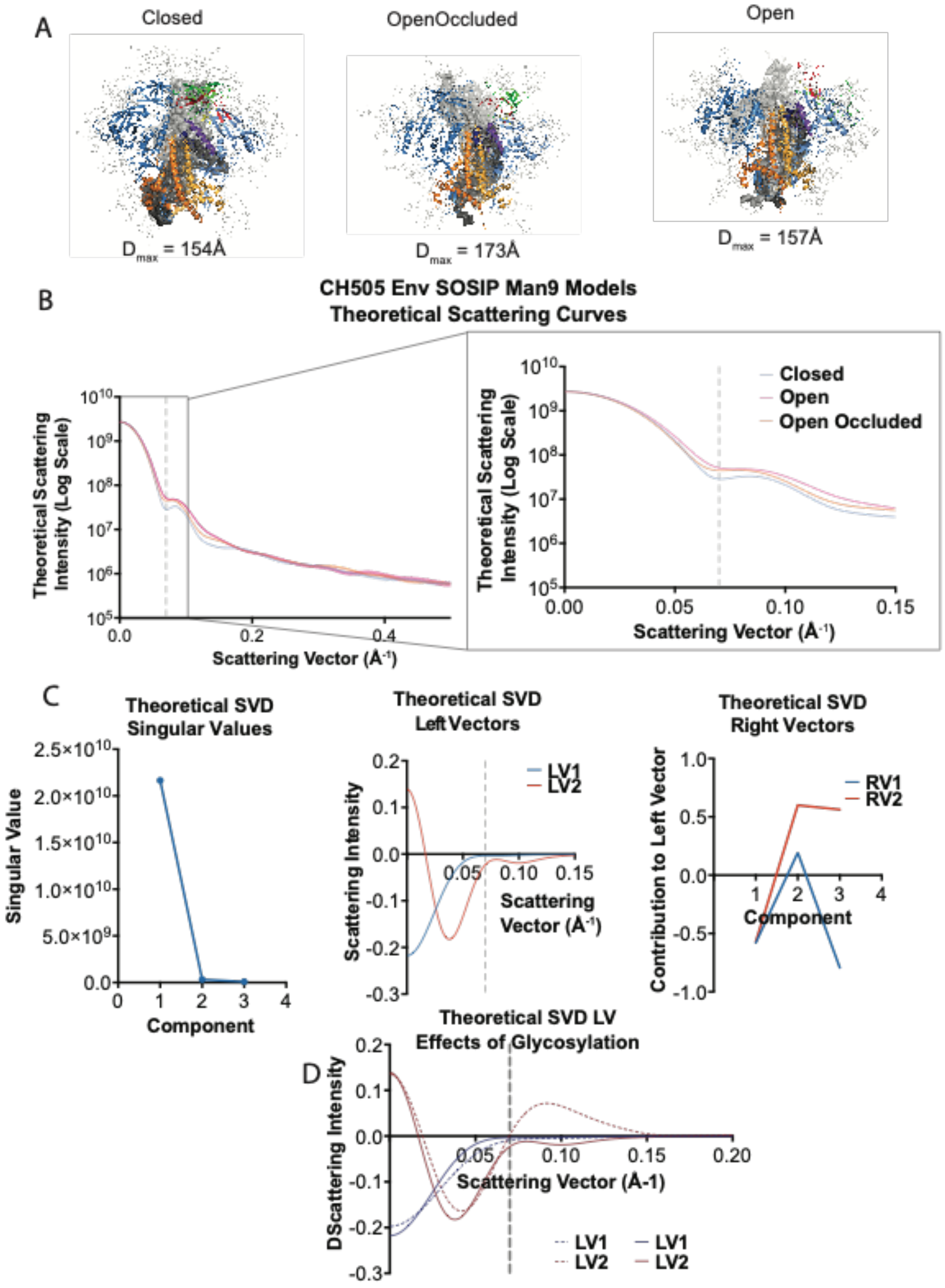

**Supplemental Figure 6: CH505TF Env SOSIP Model Glycosylated with Mannose-9. (A)** All-atom models of CH505TF Env SOSIP glycosylated with Man9 in the closed (left), Open-Occluded (*middle*), and Open conformations (*right*). **(B)** The theoretical SAXS scattering profiles showing the scattering intensity in the log scale as a function of the scattering vector in $Å^{-1}$ for Man9 glycosylated 3Closed (blue), OpenOccluded (orange), and Open (pink) CH505TF Env SOSIP models. The box shows the region detailed in the inset and the grey dash line depicts the feature peak. The standard errors of the arithmetic mean are shown as shaded regions in the plots. **(C)** The singular values of singular value decomposition (SVD) on CH505TF Env SOSIP TR, T-Jump difference profiles plotted as a function of the extracted SVD vector (*left*). SVD left vectors 1 (LV1, blue) and 2 (LV2, red) (*middle*). Dashed grey line at q=0.07 $Å^{-1}$ indicates the location of the feature peak. The SVD right vectors 1 (RV1, blue) and 2 (RV2, red) showing the contribution of LV1 and LV2 at each time delay (*right*). **(D)** SVD first left vectors (LV1, blue) and second left vectors (LV2, red) for Man9 glycosylated (solid) and non-glycosylated (dashed) CH505 Env SOSIP models.

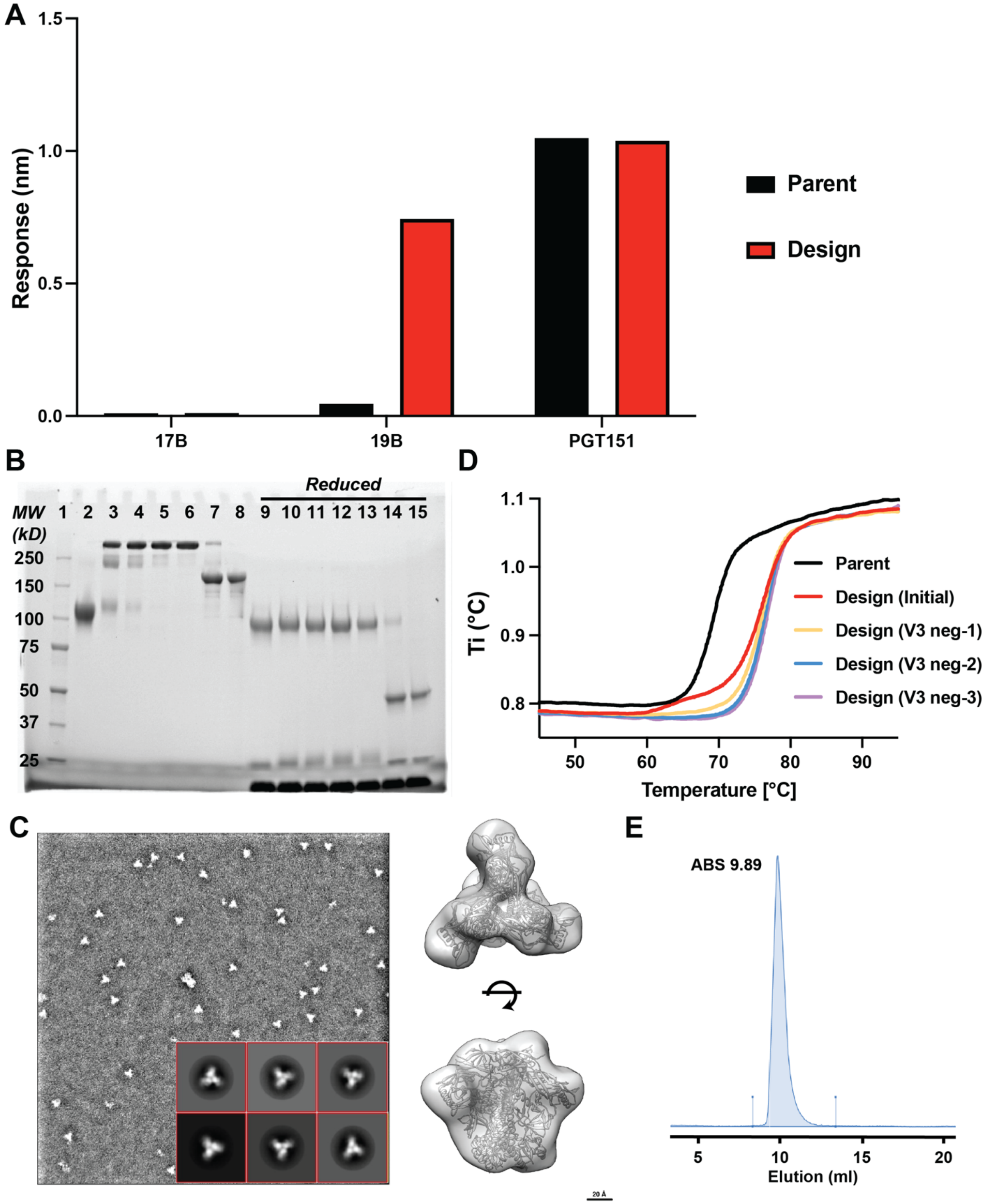

**Supplemental Figure 7: (A)** Biolayer interferometry binding responses for 17b, 19b, and PGT151 mAbs interacting with the parent and apex stapled design Env SOSIPs. **(B)** SDS-PAGE gel. Lanes are as follows: 1- Molecular weight protein marker, 2- Parent SOSIP, 3-

PGT151 purified CH505 design prior to V3 interactive 3074 based negative selection, 4-6- Sequential rounds of negative selection, 7 & 8- Eluant from Protein A column post-V3 negative selection, 9-15- Reduced samples from 2-8. **(C)** Representative NSEM micrograph, 2D-class averages, and 3D reconstruction. **(D)** Differential scanning fluorescence-based thermal melting profiles for the parent CH505 SOSIP and the apex stapled CH505 SOSIP design pre- and post-V3 negative selection iterations. **(E)** Analytical SEC profile for the final round of V3 negative selection.

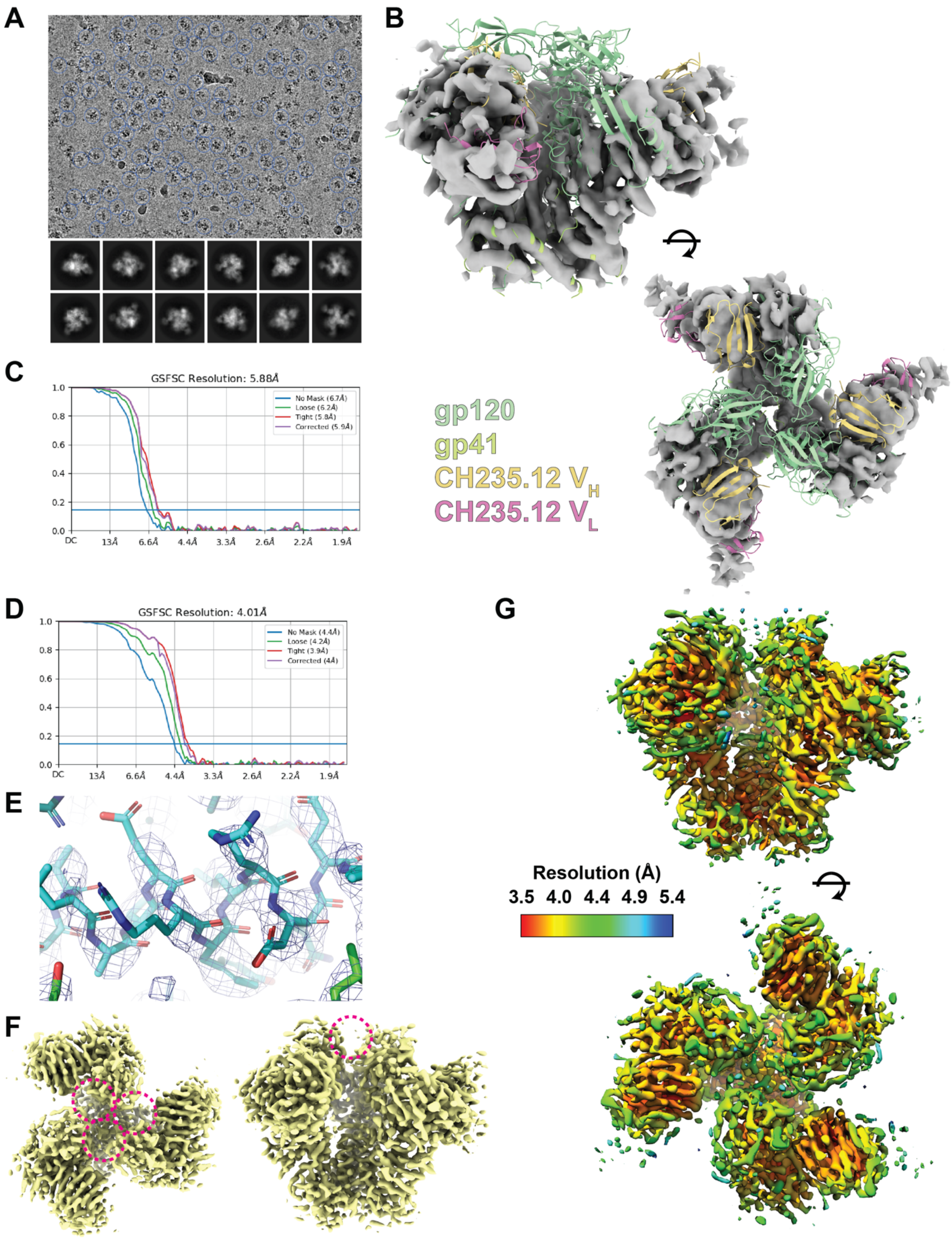

**Supplemental Figure 8: (A)** Representative cryo-EM micrograph and 2D-class averages for the apex stapled CH505 design bound to the CH235.12 Fab. **(B)** Open state map aligned with a closed state CH505 SOSIP model bound to CH235.12 Fab. **(C)** FSC plot for the Open state map. **(D)** FSC plot for the closed state map. **(E)** Representative map-to-model fit for the closed state structure. **(F)** Closed state map highlighting weak density at the trimer apex. **(G)** Local resolution map for the closed state map.

A

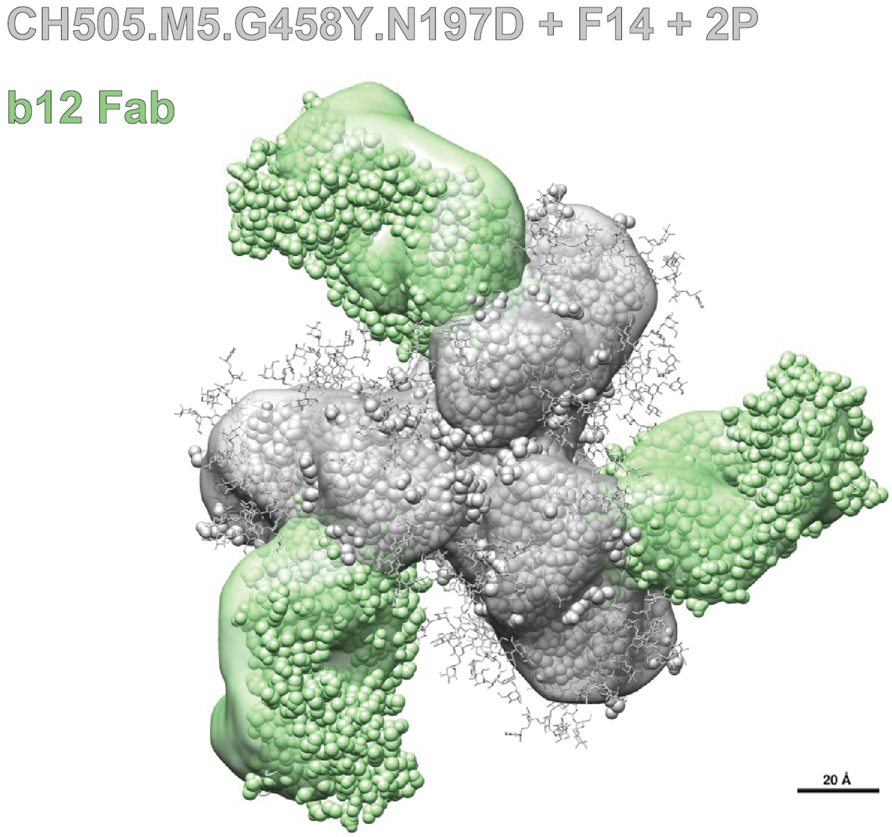
CH505.M5.G458Y.N197D + F14 + 2P
b12 Fab
20 Å

B

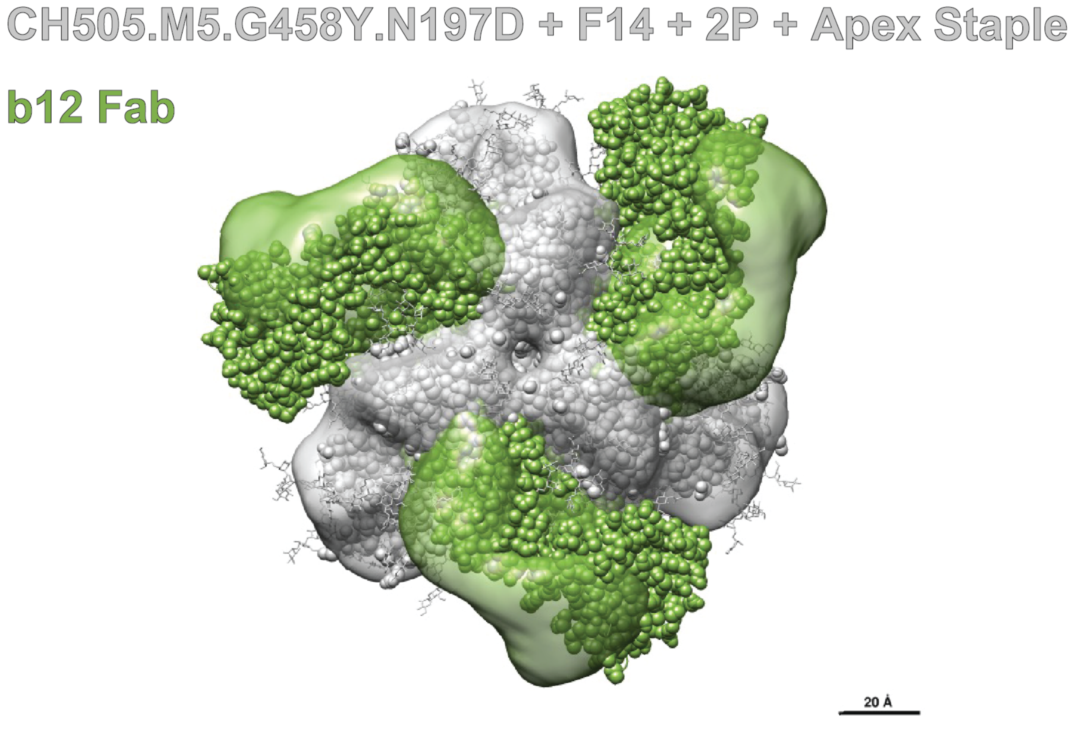
CH505.M5.G458Y.N197D + F14 + 2P + Apex Staple
b12 Fab
20 Å

**Supplemental Figure 9: (A)** Negative state 3D map and model fit of the non-apex stapled parent SOSIP (grey) bound to the b12 Fab (green) (**B)** Negative state 3D map and model fit of the apex stapled design SOSIP (grey) bound to the b12 Fab (green)

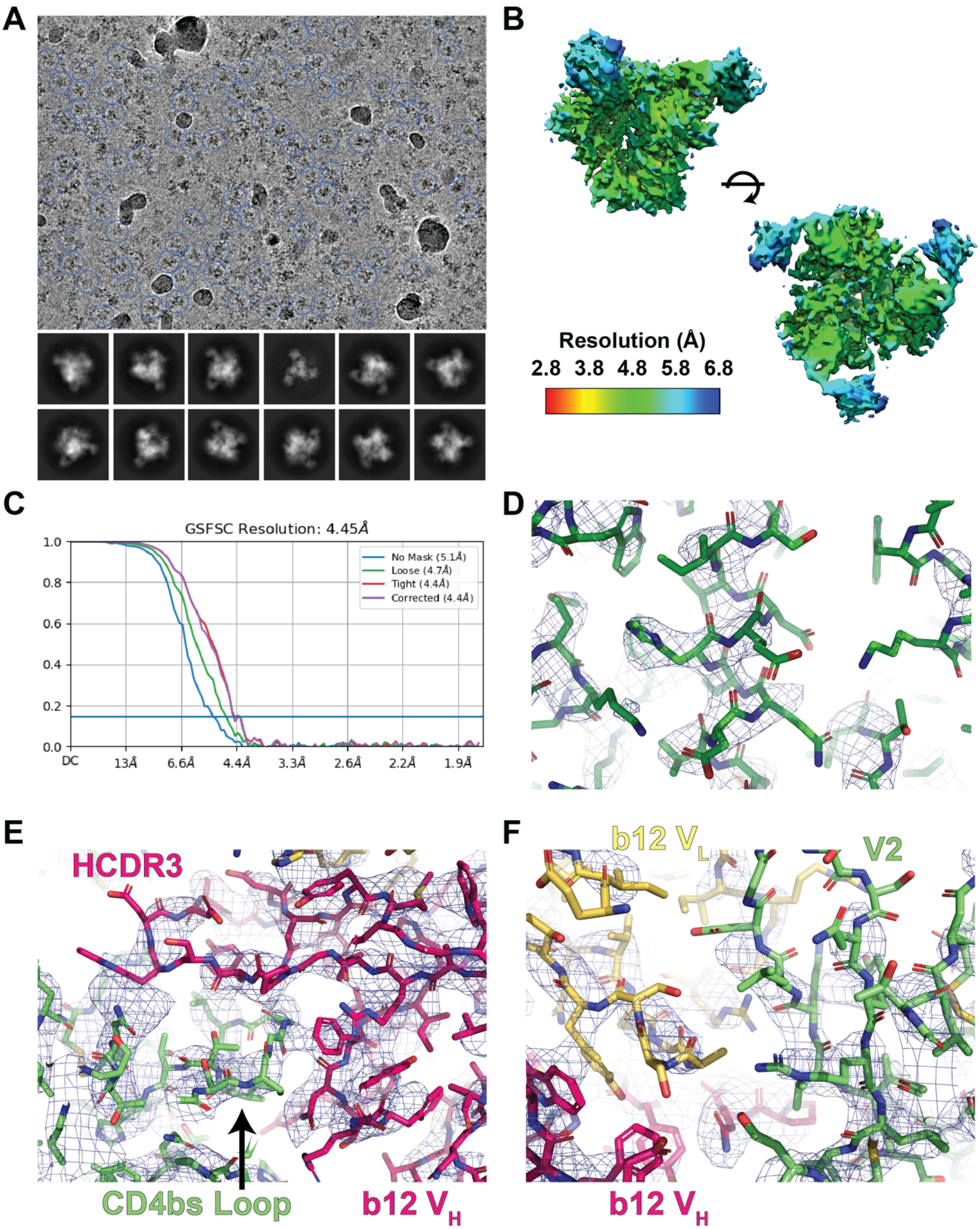

**Supplemental Figure 10: (A)** Representative cryo-EM micrograph and 2D-class averages for the apex stapled CH505 design bound to the b12 Fab. **(B)** Local resolution map. **(C)** FSC plot for the b12 bound map. **(D)** Representative map-to-model fit. **(E)** Representative map-to-model fit in the region of the b12 HCDR3 to gp120 CD4bs contact. **(F)** Representative map-to-model fit in the region of the b12 $V_L$ to gp120 V2 loop contact

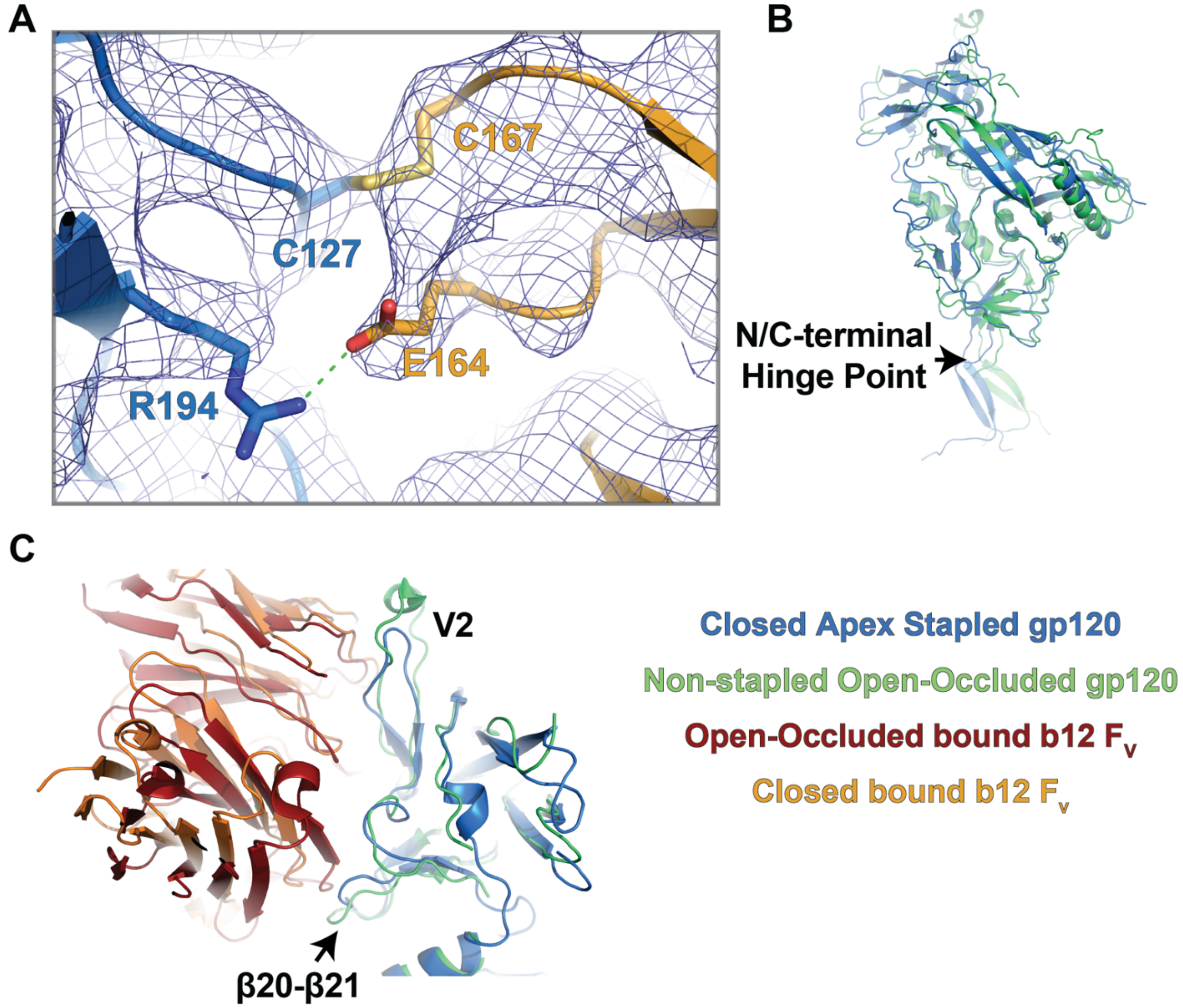

**Supplemental Figure 11: (A)** Map to model fit depicting the introduced inter-protomer apex staple. **(B and C)** Alignment of the b12 bound apex stapled CH505 SOSIP design and B41 isolate gp120 domains highlighting differences and similarities of *(B)* gp120 domains *(C)* and β20-β21 and V2 loops.

#### Supplemental Table 1: Cryo-EM Data Collection and Refinement Statistics

| | | |
|---|---|---|
| | CH505 Design + CH235.12 Fab | CH505 Design + b12 Fab |
| **Data Collection** | | |
| Microscope | Talos Arctica G3 | Talos Arctica G3 |
| Voltage (kV) | 200 | 200 |
| Electron dose ($e^-/Å^2$) | 57 | 58 |
| Detector | Gatan K3 | Gatan K3 |
| Pixel Size (Å) | 0.874 | 0.874 |
| Defocus Range (µm) | 0.1-2.88 | 0.1-2.48 |
| Magnification | 45,000 | 45,000 |
| **Reconstruction** | | |
| Software | cryoSPARC | cryoSPARC |
| Particles | 116,771 | 151,789 |
| Symmetry | C3 | C3 |
| Box size (pix) | 300 | 300 |
| Resolution (Å) (FSC0.143)$ | 4.2 | 4.5 |
| **Validation** | | |
| Protein residues | 2415 | 2439 |
| EMRinger Score | 2.5 | 1.24 |
| R.m.s. deviations | | |
| Bond lengths (Å) | 0.005 | 0.006 |
| Bond angles (°) | 0.924 | 0.998 |
| Molprobity score | 1.49 | 1.72 |
| Clash score | 2.98 | 5.43 |
| Rotamer outliers (%) | 0.05 | 0.05 |
| Ramachandran | | |
| Favored regions (%) | 94.06 | 93.38 |
| Disallowed regions (%) | 0.34 | 0.38 |

$ Resolutions are reported according to the FSC 0.143 gold-standard criterion
### Statistics are reported for the protein residues within the complex excluding the antibody constant domains
